# KOLF2.1J iTF-Microglia: A standardized platform to study microglial transcriptional regulatory networks in CNS disease

**DOI:** 10.1101/2025.05.30.657077

**Authors:** Ivan Rodriguez-Nunez, Samuel C. Bartley, Jared W. Taylor, S. Quinn Johnston, Brianne B. Rogers, Sarah K. Meadows, Kimberly M. Newberry, J. Nicholas Cochran, Richard M. Myers

## Abstract

Understanding transcriptional regulatory networks (TRNs) in microglia is key to uncovering mechanisms driving central nervous system (CNS) disorders. Human iPSC-derived models offer a tractable system for studying microglia, yet variability between lines has limited reproducibility. Here, we use the standardized KOLF2.1J iTF line to rapidly generate microglia-like cells (iTF-Microglia) and profile TRNs under homeostatic and inflammatory conditions. iTF-Microglia closely resemble primary brain microglia at both transcriptomic and epigenomic levels. Integrative analyses reveal microglia-enriched candidate *cis*-regulatory elements (cCREs) and dynamic enhancer remodeling upon differentiation and LPS+IFNG stimulation, involving key transcription factors (TFs) including NF-κB, IRF, and STAT families. TRNs active in iTF-Microglia are enriched for genetic variants linked to Alzheimer’s disease and other CNS disorders. These findings establish KOLF2.1J iTF-Microglia as a reproducible and genetically tractable platform for studying human microglial gene regulation and provide mechanistic insight into how TRN remodeling may contribute to CNS disease risk.

## Introduction

Microglia, the primary tissue-resident myeloid cells of the central nervous system (CNS), play essential roles in development, homeostasis, immune surveillance, and inflammation. Dysregulated microglial function is implicated in the onset and progression of various brain disorders (Gao et al., 2023; Prinz and Priller, 2014), and genetic risk variants for neurodegenerative diseases are often enriched in microglial transcriptional networks (Askarova et al., 2024). However, single-cell/nuclei RNA-seq and ATAC-seq studies have underscored the transcriptional and epigenetic heterogeneity and plasticity of microglia, complicating efforts to interpret the impact of disease-risk variants (Adams et al., 2024; Anderson et al., 2023; Gamache et al., 2023; Lee et al., 2023; Morabito et al., 2021; Prater et al., 2023; Shi et al., 2024; Sun et al., 2023; Xiong et al., 2023; Zhao et al., 2024).

Studying microglia is challenging due to their inaccessibility, reliance on environmental cues for maintaining tissue-specific identity, relative sparsity compared to other cell types, and interspecies differences (Drummond and Wisniewski, 2017; Gosselin et al., 2017; Li et al., 2023). Human pluripotent stem cells (PSCs) have enabled the generation of microglia-like cells (shortened to “microglia” in subsequent text) through protocols that recapitulate primitive hematopoiesis with maturation in monoculture or co-culture systems (Abud et al., 2017; Douvaras et al., 2017; Guttikonda et al., 2021; Konttinen et al., 2019; McQuade et al., 2018; Muffat et al., 2016; Washer et al., 2022). A recent approach uses a doxycycline-inducible system to drive differentiation via six transcription factors (TFs): IRF5, IRF8, CEBPA, CEBPB, PU.1, and MAFB, with resultant cells denoted as iTF-Microglia (Dräger et al., 2022). This rapid protocol produces functional microglia capable of synaptosome phagocytosis, inflammatory responses, and integration into co-culture systems with induced PSCs (iPSC)-derived neurons.

PSC-derived microglia offer key advantages, including scalability, genetic manipulation, and relevance to human biology, but reproducibility is often affected by donor-to-donor genetic variation (Volpato and Webber, 2020). The iPSC Neurodegenerative Disease Initiative (iNDI) addressed this by engineering the KOLF2.1J iPSC line, a genomically stable reference line neutral to polygenic Alzheimer’s disease (AD) risk (Pantazis et al., 2022). Because of the particular scrutiny of this line, it was found that heterozygous copy number variants (CNVs) are associated with neurodevelopmental traits (Gracia-Diaz et al., 2024), however, recent analyses have proven that KOLF2.1J is suitable for neurological disease modeling (Dobner et al., 2024a; Dou et al., 2025; Martinez-Sanchez et al., 2025; Nam and Ordureau, 2022; Ryan et al., 2024). Additionally, the availability of KOLF2.1J isogenic lines carrying AD and related dementias (ADRD) variants (Ramos et al., 2021) offer a valuable platform for genotype-to-phenotype studies (Fang et al., 2025).

Here, we used a KOLF2.1J iPSC line carrying the inducible iTF cassette (KOLF2.1J iTF), as described by Dräger et al. (Dräger et al., 2022). We leveraged this model to investigate transcriptional regulatory networks (TRNs) involved in microglial differentiation and response to lipopolysaccharide (LPS) and interferon-gamma (IFNG). Specifically, we identified candidate *cis*-regulatory elements (cCREs), upstream TFs and target genes, and evaluated the enrichment of neurodegenerative risk variants in these networks, offering mechanistic insights into microglial contributions to CNS disease.

## Results

### KOLF2.1J iTF-Microglia transcriptome resembles other in vitro microglia models and brain microglia

Differentiation of KOLF2.1J iTF-iPSCs into iTF-Microglia induced widespread transcriptional changes, including upregulation of core homeostatic microglia markers such as *CX3CR1*, *P2RY12*, *TMEM119*, *TGFBR1*, *GPR34*, *MERTK*, *PROS1*, and *HEXB* (Gerrits et al., 2020) (**Figure 1A; see also Table S1**). *SALL1*, a known homeostatic microglia marker and critical TF for microglia identity (Buttgereit et al., 2016; Fixsen et al., 2023), was more expressed in iTF-iPSCs. Notably, SALL1 is also a potent activator of iPSC reprogramming in cooperation with NANOG (Yang et al., 2019).

**Figure 1.**
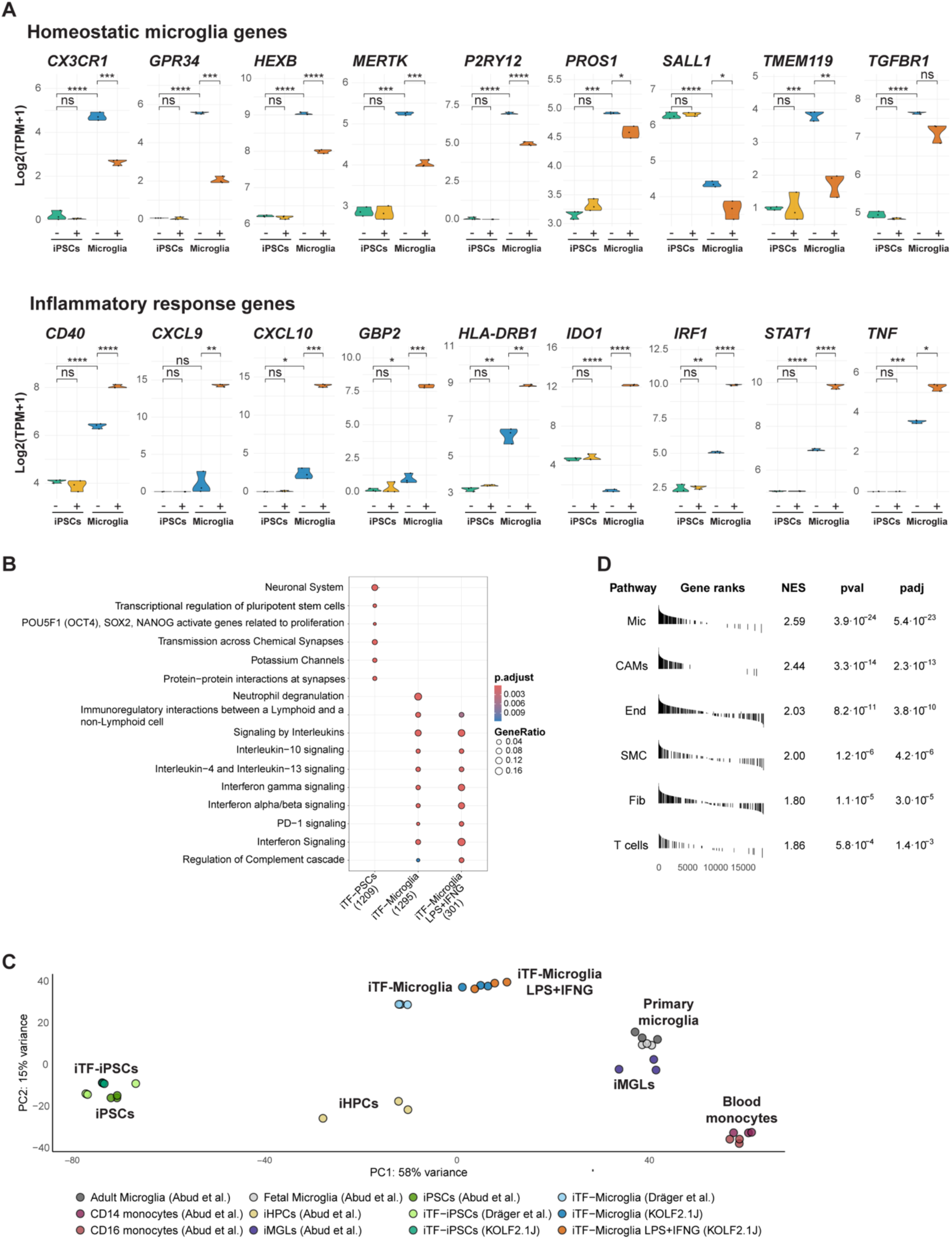
Transcriptomic characterization of KOLF2.1J iTF-Microglia. (**A**) Expression of homeostatic microglia markers and inflammatory response genes in untreated (−) and LPS+IFNG-treated (+) iTF-Microglia and iTF-iPSCs. Violin plots display gene expression distributions; individual dots represent expression values from each replicate (n = 3). Significance was determined using pairwise t-tests (ns: p > 0.5, * p ≤ 0.05, ** p ≤ 0.01, *** p ≤ 0.001, **** p ≤ 0.0001). (**B**) Reactome pathway enrichment of DEGs (LFC > 2, padj < 0.05) from three comparisons: iTF-iPSCs up (vs. iTF-Microglia), iTF-Microglia up (vs. iTF-iPSCs), and iTF-Microglia LPS+IFNG up (vs. iTF-Microglia). Pathways were identified using compareCluster (pvalueCutoff = 0.05) and plotted with showCategory = 6. See also Table S2. (**C**) PCA of KOLF2.1J iTF cells alongside other iPSCs, iHPCs, in vitro-derived microglia, primary adult and fetal microglia, and blood monocytes. (**D**) GSEA using DESeq2-ranked genes and brain cell-type markers. Enriched cell types include CNS-associated macrophages (CAMs), Endothelial cells (End), fibroblasts (Fib), microglia (Mic), smooth muscle cells (SMC), and T cells.

LPS+IFNG treatment in iTF-Microglia downregulated homeostatic genes and upregulated immune response genes, such as *CXCL9*, *CXCL10*, and *TNF* (involved in immune cell recruitment and neuroinflammation); *STAT1* and *IRF1* (key IFNG signaling mediators); *HLA-DRB1* and *CD4* (antigen presentation); and *GBP2* (an immune defense protein induced by IFNG) (Carter et al., 2007; McLellan et al., 1999; Rauf et al., 2022; Schoenborn and Wilson, 2007; You et al., 2024). *IDO1* that encodes a metabolic enzyme linked to IFNG responses and neurodegeneration (Pallotta et al., 2022) was also upregulated (**Figure 1A; see also Table S1**). Notably, *IDO1* expression was higher in iTF-iPSCs than in untreated iTF-Microglia, consistent with its role as a marker of iPSC contamination and maintaining pluripotency of embryonic stem cells (ESCs) via glycolysis (Lemmens et al., 2023; Liu et al., 2019). In contrast, LPS+IFNG had minimal transcriptional impact on iTF-iPSCs (**Figure S1A; see also Table S1**), likely due to low or absent expression of IFNG pathway mediators and LPS receptors (**Figure S1B**).

Pathway analysis of differentially expressed genes (DEGs) (LFC > 2, padj < 0.05) showed enrichment of immune-related pathways in iTF-Microglia, including IFNG signaling, interleukin signaling and complement regulation. These pathways were further enhanced upon LPS+IFNG treatment. In contrast, iPSCs showed enrichment for pluripotency-associated pathways (**Figure 1B; see also Table S2**).

Principal component analysis (PCA) showed iTF-Microglia clustering together, regardless of WTC11 (from Dräger et al., 2022) or KOLF2.1J background or LPS+IFNG treatment. iPSCs formed a separate cluster. Induced microglia-like cells (iMGLs), differentiated for over 30 days using growth factors (Abud et al., 2017), clustered closest to primary microglia, while induced hematopoietic progenitor cells (iHPCs), an intermediate stage in the Abud et al. differentiation protocol, and blood monocytes formed distinct, separate clusters (**Figure 1C**). Hierarchical clustering based on cell-type marker genes confirmed these groupings; iPSCs and iHPCs formed their own subclusters, monocytes clustered separately, and all microglial models grouped together, with iMGLs showing the highest similarity to primary microglia (**Figure S1C**).

Gene set enrichment analysis (GSEA) revealed that genes upregulated in iTF-Microglia relative to iTF-iPSCs were enriched for marker genes of brain-resident cell types (Mathys et al., 2024), especially microglia and CNS-associated macrophages (**Figure 1D**). In contrast, genes upregulated in iTF-iPSCs were not enriched for any brain cell type. Within iTF-Microglia, LPS+IFNG treatment upregulated gene signatures associated with interferon, inflammation and reactive microglial states, while untreated cells were mainly enriched for proliferative, enhanced-redox, and surveilling microglial subtypes (Green et al., 2024) (**Figure S1D**).

### KOLF2.1J iTF-Microglia cCREs are enriched for immune response pathways

The co-occurrence of chromatin accessibility and H3K27ac enrichment marks active enhancers and promoters, which can be used to identify cCREs relevant to gene regulation (Stępniak et al., 2021). Using this approach, we identified 37,193 cCREs in iTF-iPSCs, 47,315 in iTF-Microglia, and 46,795 in LPS+IFNG-treated iTF-Microglia. Many were cell type- or state-specific, with most overlapping distal enhancer-like sequences (dELS) from ENCODE4 (**Figure 2A**). In contrast, cCREs shared across iTF-iPSCs, iTF-Microglia, and LPS+IFNG-treated iTF-Microglia were enriched for promoter-like sequences (PLS) and proximal enhancer-like sequences (pELS).

**Figure 2.**
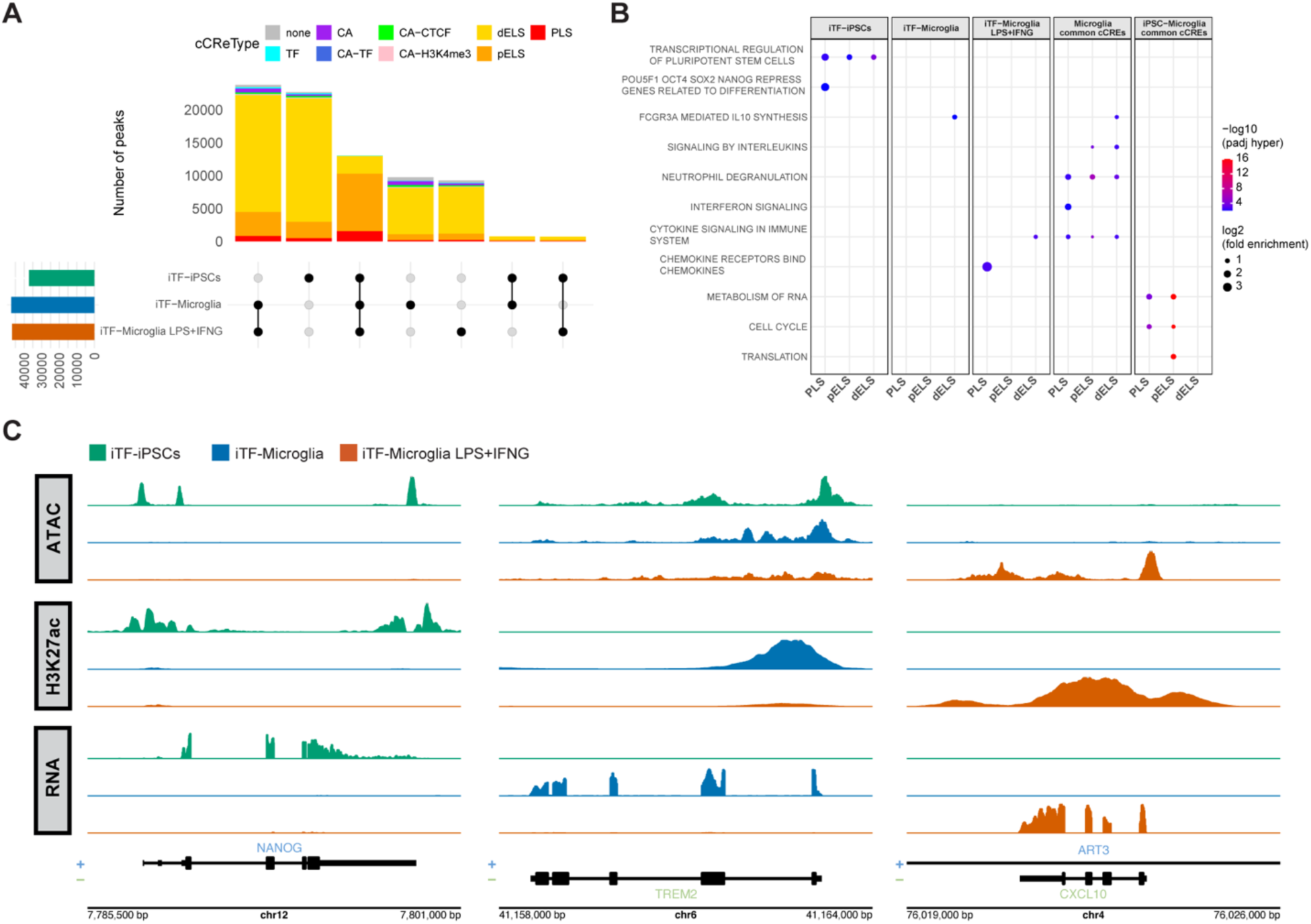
KOLF2.1J iTF-Microglia cCREs are enriched for immune response pathways. **(A)** Overlap of cCREs identified in iTF-iPSCs, untreated iTF-Microglia, and LPS+IFNG-treated iTF-Microglia, annotated using ENCODE4 categories: CA (chromatin accessibility), CA-CTCF (CA with CTCF binding), dELS (distal enhancer-like sequence), PLS (promoter-like sequence), TF (transcription factor binding), CA-TF (CA with TF binding), CA-H3K4me3 (CA with H3K4me3), and pELS (proximal enhancer-like sequence). (**B**) Reactome pathway enrichment of cCREs from each condition. Pathways with adjusted hypergeometric p <= 0.01 were retained and manually curated. See also Table S3. (**C**) Representative genome browser tracks showing ATAC-seq, H3K27ac, and RNA-seq signal at *NANOG*, *TREM2*, and *CXCL10* loci.

Functional enrichment analysis revealed that cCREs in iTF-iPSCs were associated with pluripotency pathways, while cCREs shared by untreated and LPS+IFNG-treated iTF-Microglia were enriched for immune pathways. LPS+IFNG-specific cCREs were particularly enriched for chemokine signaling. cCREs common to iTF-iPSCs, untreated iTF-Microglia and LPS+IFNG-treated iTF-Microglia were linked to core cellular functions, such as RNA processing, protein translation, and cell cycle regulation (**Figure 2B; see also Table S3**).

Among the identified cCREs, we found overlapping ATAC-seq and H3K27ac peaks at both ends of the *NANOG* gene in iTF-iPSCs, a TF used as a pluripotency marker (Dobner et al., 2024b). In contrast, *TREM2*, a gene expressed in microglia and implicated in AD and Nasu-Hakola disease pathogenesis (Yeh et al., 2017), showed similar chromatin accessibility across iTF-iPSCs, untreated iTF-Microglia, and LPS+IFNG-treated iTF-Microglia, while an H3K27ac peak was only present in untreated iTF-Microglia. Loss of this peak in LPS+IFNG-treated iTF-Microglia correlated with strong *TREM2* downregulation. In contrast, the early-response IFNG gene *CXCL10* (Carter et al., 2007) showed an active cCRE signature exclusively in LPS+IFNG-treated iTF-Microglia (**Figure 2C**).

### KOLF2.1J iTF-Microglia TFs regulate microglial identity and response

TF binding at CREs plays a central role in regulating gene expression. Lineage-determining TFs are essential for initiating and maintaining open chromatin states associated with cell identity, while signal-dependent TFs adapt gene expression in response to external cues (Field and Adelman, 2020). In macrophages, inflammatory responses are marked by changes in H3K27ac levels at enhancers, which are enriched for binding motifs of TFs involved in immune signaling (Herrera-Uribe et al., 2020). Notably, H3K27ac has also been implicated in microglial immune memory following LPS exposure (Huang et al., 2023).

We found that LPS+IFNG-treated iTF-Microglia increased chromatin accessibility and H3K27ac marking at numerous cCREs. Only one cCRE showed reduced accessibility compared to untreated iTF-Microglia, while approximately 400 cCREs showed decreased H3K27ac signal (**Figures S2A and S2B**), suggesting that H3K27ac loss contributes to transcriptional repression during LPS+IFNG treatment.

To identify candidate TFs driving microglial identity and inflammatory responses, we applied randomized lasso stability selection approach using the monaLisa package (Machlab et al., 2022). This uncovered TF motifs associated with chromatin accessibility changes during differentiation (iTF-iPSCs vs. iTF-Microglia) and H3K27ac remodeling upon stimulation (LPS+IFNG-treated iTF-Microglia vs. iTF-Microglia).

cCREs more accessible in iTF-iPSCs were enriched for motifs of SOX and POU TF families, key regulators of pluripotency, self-renewal, and reprogramming (Abdelalim et al., 2014; Yusa et al., 2009), as well as TEAD motifs, known to enhance iPSC reprogramming efficiency (Huang et al., 2021). In contrast, cCREs active in iTF-Microglia were enriched for ETS and C/EBP motifs, including PU.1 (*SPI1*) and CEBPA, lineage-determining TFs essential for microglia development (Chen et al., 2021) and both encoded by the iTF cassette inserted into KOLF2.1J cells (**Figure 3A**).

**Figure 3.**
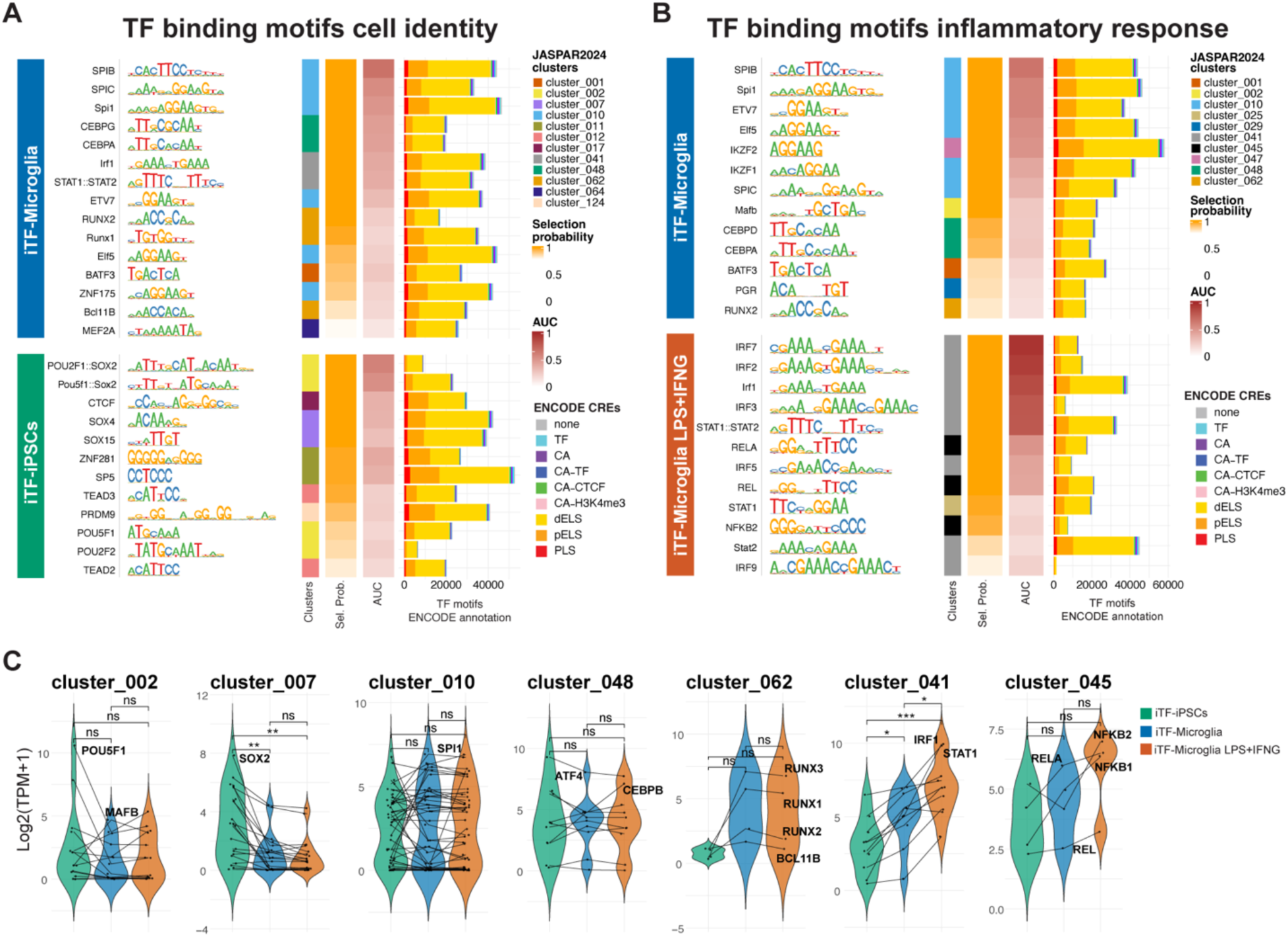
TF motifs linked to microglial differentiation and inflammatory response. **(A)** TF motifs enriched in cCREs with different chromatin accessibility between iTF-iPSCs and iTF-Microglia. Motifs are ranked by selection probability and AUC from randomized lasso stability selection; corresponding ENCODE annotations are shown. (**B**) TF motifs enriched in cCREs with different H3K27ac signal between LPS+IFNG-treated and untreated iTF-Microglia. (**C**) Expression levels of TFs associated with each JASPAR2024 motif cluster identified in panels A and B. Violin shapes indicate the overall distribution of TF expression within each cluster and condition. Each dot represents the average expression (from n = 3 replicates) of an individual TF. Lines connect the same TF across conditions. Statistical significance between conditions was assessed using two-sided t-tests (ns, p > 0.05; *p ≤ 0.05; **p ≤ 0.01; ***p ≤ 0.001).

Comparison of LPS+IFNG-treated versus untreated iTF-Microglia revealed that cCREs with higher H3K27ac in the untreated state were enriched for motifs of homeostatic TFs such as RUNX2, MAFB, and IKZF1, which are implicated in maintaining microglial identity and modulating neurodegeneration (Ballasch et al., 2023; Gosselin et al., 2017; Prater et al., 2023; Yamasaki et al., 2024). PU.1 (*SPI1*) and CEBPA motifs were also found, consistent with their dual roles in homeostasis and inflammation (Gao et al., 2019; Yeh and Ikezu, 2019; Zhang et al., 2024). In contrast, cCREs with increased H3K27ac upon LPS+IFNG treatment were enriched for NF-κB, IRFs, and STATs motifs, signal-dependent TFs that mediate of microglial inflammatory responses (Papageorgiou et al., 2016; Wang et al., 2024; Young and Denovan-Wright, 2024) (**Figure 3B**). Since lasso regression can arbitrarily select only a subset of correlated motifs within motif clusters, we also examined the expression of TFs corresponding to the thirteen JASPAR2024 motif clusters identified by monaLisa. For example, ELF5 was selected from Cluster 10 (enriched in untreated iTF-Microglia) but had low expression in iTF-iPSCs and iTF-Microglia. In contrast, ELF1 and ELK3, also in Cluster 10, were highly expressed in iTF-Microglia and have known roles in microglial development (Kracht et al., 2020). In cluster 47, IKZF2 was unchanged between iTF-iPSCs and iTF-Microglia, while PRDM1, a co-cluster member, was more expressed in iTF-Microglia, aligning with its known enrichment in adult microglia (Yaqubi et al., 2023). Cluster 7 SOX TFs (enriched in iTF-iPSCs cCREs) showed higher expression in iTF-iPSCs, while cluster 41 TFs (enriched in LPS+IFNG-stimulated cCREs), were more upregulated under inflammatory conditions (**Figure 3C**).

### KOLF2.1J iTF-Microglia TRNs target genes

Assigning distal CREs to their target genes is a critical yet complex task for understanding gene regulation. While the most common approach assigns enhancers to the closest gene (Kaikkonen et al., 2013), many enhancers often bypass the nearest gene to regulate more distal ones (Yao et al., 2015).

To improve confidence in cCRE-to-gene associations, we applied different methods: (1) closest gene, (2) DegCre, (3) activity-by-contact (ABC) model, and (4) brain microglia links. Collectively, these methods identified numerous cCRE-to-gene associations (**Table 1**).

**Table 1.** Number of cCRE-to-gene associations for each mapping method.

|  |  | **cCREs** | **Target genes** |
| --- | --- | --- | --- |
| **closest**<br>**TSS** | *iTF-iPSCs* | 37,193 | 13,295 |
|  | *iTF-Microglia* | 47,315 | 14,472 |
|  | *iTF-Microglia LPS+IFNG* | 46,795 | 14,426 |
| **DegCre** | *iTF-iPSCs* | 28,393 | 6,806 |
|  | *iTF-Microglia differentiated* | 21,392 | 4,083 |
|  | *iTF-Microglia untreated* | 27,962 | 5,122 |
|  | *iTF-Microglia LPS+IFNG* | 16,238 | 4,059 |
| **ABC** | *iTF-iPSCs* | 19,963 | 12,024 |
|  | *iTF-Microglia* | 23,549 | 11,439 |
|  | *iTF-Microglia LPS+IFNG* | 23,929 | 11,286 |
| **Brain**<br>**microglia link** | *iTF-iPSCs* | 10,424 | 9,879 |
|  | *iTF-Microglia* | 20,424 | 11,475 |
|  | *iTF-Microglia LPS+IFNG* | 19,756 | 11,373 |

Notably, both untreated and LPS+IFNG-treated iTF-Microglia showed significantly greater overlap with brain microglia cCREs (from Anderson et al., 2023) than iTF-iPSCs (Fisher’s exact test, p < 2.2 × 10^−16^ for both comparisons). The odds ratios were 1.95 and 1.88, respectively, indicating nearly double the likelihood of overlap. Additionally, untreated iTF-Microglia exhibited a modest but significant enrichment over LPS+IFNG-treated iTF-Microglia (odds ratio = 1.04, p = 0.0034). For DegCre, we used differential chromatin accessibility to map cCRE-to-gene associations involved in microglial differentiation (iTF-iPSCs vs. iTF-Microglia), and H3K27ac changes to infer inflammatory TRNs (LPS+IFNG-treated iTF-Microglia vs. untreated iTF-Microglia).

The distribution of the distances between cCREs and their predicted target genes peaked at TSS and distal regions around 100 kb from TSS (**Figure S3A**).

Most cCRE-to-gene associations were method-specific, with limited overlap across models (**Figure 4A; see also Figure S3B**). The ABC model and brain microglia links tended to assign promoter-associated cCREs (defined as regions within −2 kb to +0.2 kb of GENCODE v46 TSS), likely due to high ATAC (and H3K27ac for ABC) signal at promoters in gene-dense regions. This was supported by significantly higher accumulation of ATAC and H3K27ac reads at cCREs overlapping promoters compared to enhancers, which were defined as cCREs not overlapping promoters (**Figure S3C**). In contrast, DegCre identified more distal enhancer-associated cCREs by linking differential gene expression with changes in chromatin accessibility and H3K27ac, which are primarily found in distal enhancers (**Figures S2A and S2B**), making DegCre more sensitive to distal enhancer activity and offering a complementary approach for mapping cCRE-to-gene associations.

**Figure 4.**
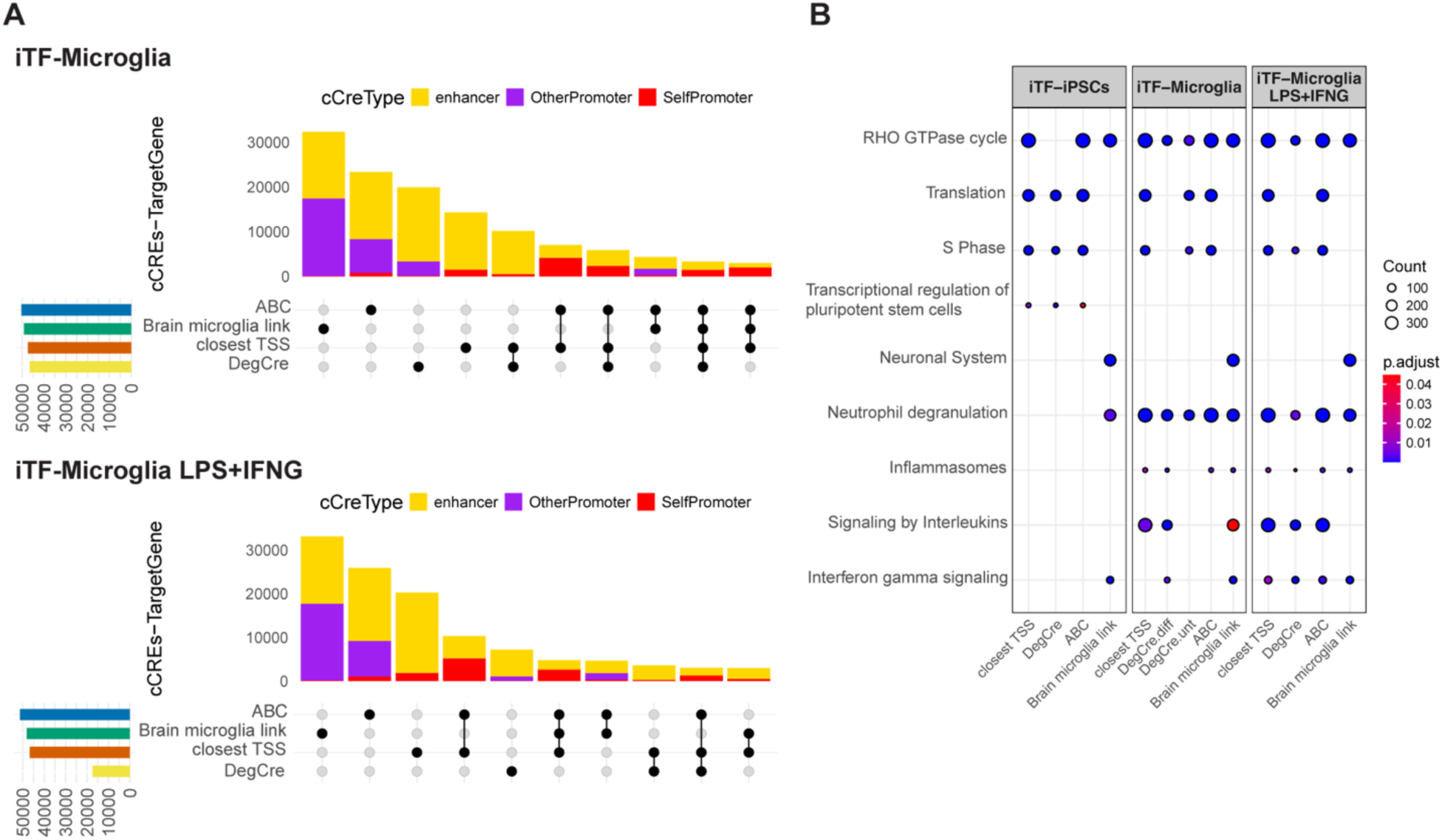
Mapping cCRE-to-gene associations in KOLF2.1J iTF-Microglia. (**A**) Overlap of cCRE-to-gene associations identified by four mapping strategies in untreated and LPS+IFNG-treated iTF-Microglia. See also Figure S3B. (**B**) Reactome pathway enrichment of predicted target genes across cell states and mapping methods. Pathways were identified using compareCluster (pvalueCutoff = 0.05) and plotted pathways were manually selected from the significantly enriched terms. See also Table S4.

Pathway enrichment analysis (**Figure 4B; see also Table S4**) revealed that cCRE-to-gene associations in iTF-iPSCs, iTF-Microglia, and LPS+IFNG-treated iTF-Microglia were enriched for protein translation and S-phase proliferation pathways, which were absent in brain microglia. Instead, brain microglia were enriched for neuronal system pathways, likely reflecting differences between in vitro culture and post-mortem brain tissue environments.

Despite these differences, several pathways were shared across conditions. The RHO GTPase cycle, which regulates cytoskeletal and signaling processes (Jaffe and Hall, 2005), was consistently enriched. Immune-related pathways showed distinct patterns, neutrophil degranulation, inflammasomes and interleukin signaling were enriched in both iTF-Microglia and brain microglia, while IFNG signaling was especially enriched in LPS+IFNG-treated iTF-Microglia and brain microglia.

These findings highlight the complexity of linking cCREs to target genes, which can be shaped by cell type, environmental context, and the cCRE-to-gene prediction method used.

### KOLF2.1J iTF-Microglia TRNs are enriched for neurodegenerative and autoimmune risk variants

GWAS have identified many genomic loci associated with disease risk, but linking these to specific variants, genes, molecular pathways, and cell types remains challenging. To evaluate disease risk variant enrichment in iTF-iPSCs and iTF-Microglia (with or without LPS+IFNG), we integrated transcriptomic and epigenomic data with GWAS summary statistics using MAGMA (De Leeuw et al., 2015) and stratified linkage disequilibrium (LD) score regression (sLDSC) (Bulik-Sullivan et al., 2015; Finucane et al., 2015).

MAGMA analysis revealed significant associations between genes expressed in untreated iTF-Microglia and AD risk variants from different GWAS. LPS+IFNG-treated iTF-Microglia showed associations with one AD dataset and variants linked to multiple sclerosis (MS), inflammatory bowel disease (IBD), and systemic lupus erythematosus (SLE) (**Figure 5A**).

**Figure 5.**
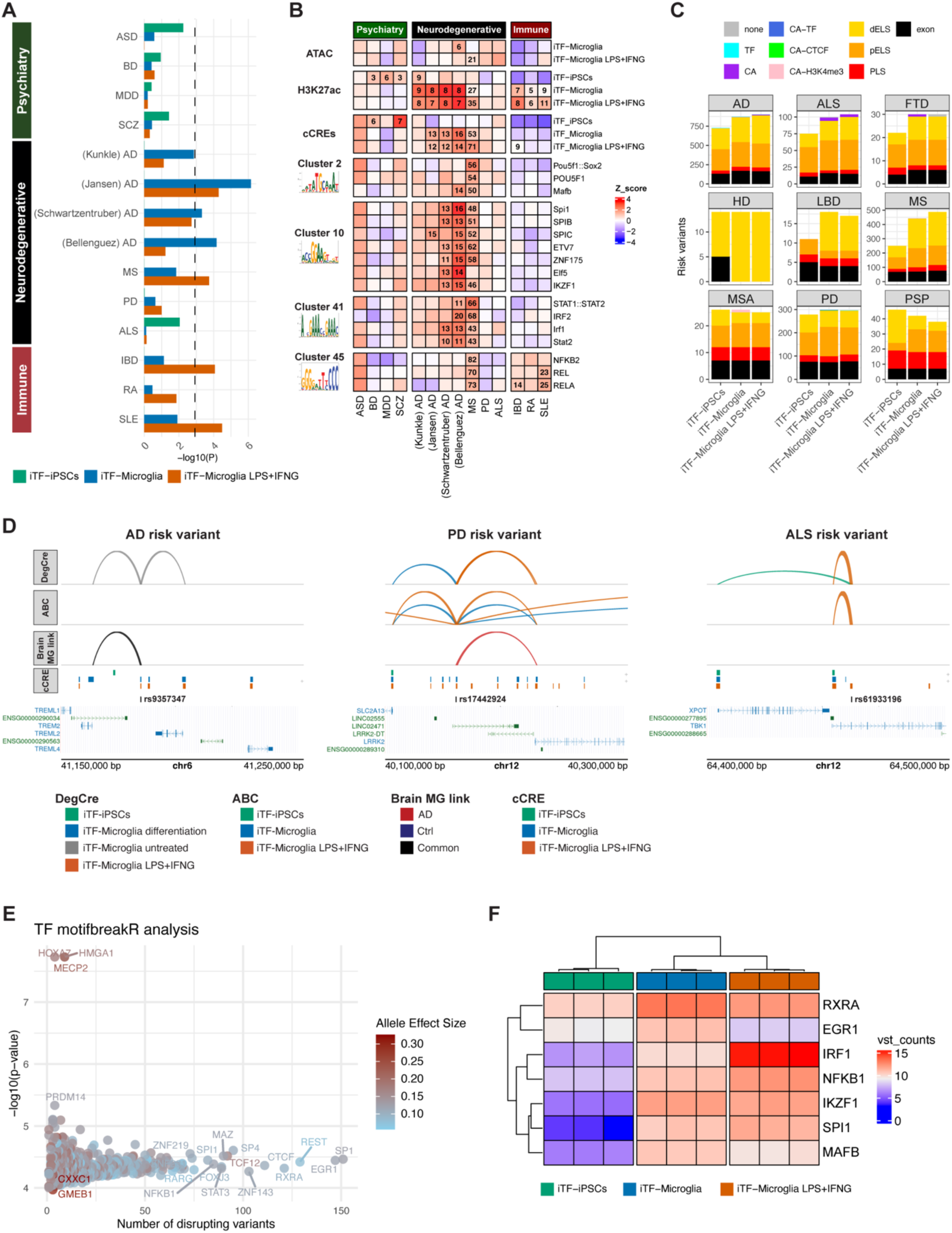
Mapping disease-risk variants to KOLF2.1J TRNs. (**A**) MAGMA analysis showing enrichment of disease-risk variants in genes expressed in iTF-iPSCs, untreated iTF-Microglia, and LPS+IFNG-treated iTF-Microglia. The black dashed line indicates a Bonferroni-corrected significance threshold of p = 0.05. (**B**) Heatmap of sLDSC coefficient Z scores for ATAC-seq peaks, H3K27ac peaks, and cCREs across conditions. Feature–disease pairs with Benjamini-Hochberg corrected p < 0.05 are annotated with their enrichment scores. See also Table S5. (**C**) Number of neurodegenerative risk variants overlapping cCREs from iTF-iPSCs, untreated iTF-Microglia, and LPS+IFNG-treated iTF-Microglia. (**D**) Linkage plots showing cCREs containing risk variants and their predicted target genes, based on DegCre, ABC, and brain microglia links models. See also Table S6. (**E**) TFs with motifs disrupted by disease-risk variants were identified using motifbreakR (strong effects only). TFs were filtered for gene expression (TPM > 1). For each TF, the lowest p-value per allele (Ref or Alt) was used, and the median –log_10_(p-value) across variants is shown. Dot color reflects median effect size. See also Table S7. (**F**) Expression levels of selected TFs whose binding motifs are disrupted by disease variants.

sLDSC analysis showed stronger enrichment in H3K27ac-marked regions than ATAC-seq peaks. Among cCREs (defined by overlapping ATAC and H3K27ac peaks), those in iTF-iPSCs were enriched for bipolar disorder (BD) and schizophrenia (SCZ) variants, while cCREs from both untreated and LPS+IFNG-treated iTF-Microglia were enriched for AD and MS variants (**Figure 5B; see also Table S5**). Disease variant enrichment was also observed in cCREs grouped by TF motifs identified through monaLisa. For example, *SPI1*-associated cluster 10 and IRF-containing cluster 41 were enriched for AD and MS, while cluster 45, marked by NF-κB motifs, was enriched for MS, IBD and SLE. Cluster 2 includes POU TFs that are enriched for MS, and MAFB, a key regulator of adult microglia gene expression and a putative driver of AD-specific CREs (Anderson et al., 2023; Yamasaki et al., 2024), that showed enrichment for both MS and AD variants.

To identify candidate disease genes, we intersected cCREs and their predicted target genes with 62,300 neurodegenerative risk variants. A large fraction of variants mapped to cCREs near gene promoters and exons (**Figure 5C**).

Key examples included variants associated with AD, Parkinson’s disease (PD) and amyotrophic lateral sclerosis (ALS) (**Figure 5D; see also Table S6**). **rs9357347 (AD)**, a variant that has been associated with *TREM2* expression and amyloid/tau pathology (Carrasquillo et al., 2017; Tian et al., 2019), was located in a cCRE identified in both untreated and LPS+IFNG-treated iTF-Microglia, and mapped to *TREM2* and *TREML2* in untreated iTF-Microglia via DegCre (H3K27ac signal and *TREM2* expression were higher in untreated iTF-Microglia compared to LPS+IFNG-treated iTF-Microglia) and brain microglia links from AD and control samples. **rs17442924 (PD)** overlapped a cCRE present in both untreated and LPS+IFNG-treated iTF-Microglia mapping to *LRRK2* via DegCre, ABC, and a brain microglia link from AD samples. Hi-C loop data supported this association. *LRRK2* mutations are known to drive microglia-mediated PD neurodegeneration (Langston et al., 2022; Panagiotakopoulou et al., 2020). **rs61933196 (ALS)** located in a cCRE specific to LPS+IFNG-treated iTF-Microglia, was mapped via DegCre and ABC to *TBK1*, a gene implicated in familial ALS and microglia activation (Brenner et al., 2019; Cirulli et al., 2015). Although also linked to *XPOT* in iTF-iPSCs via DegCre, no cCRE was detected in those cells. However, higher *XPOT* expression and chromatin accessibility in iTF-iPSCs relative to untreated iTF-Microglia suggested this association.

Additional examples listed in **Table S6** include: **rs6733839 (AD, Lewy body dementia (LBD))** and **rs72838287 (AD)**, both located within a microglia-specific enhancer at the *BIN1* locus (Nott et al., 2019). Notably, rs72838287 showed allelic differential activity in massively parallel reporter assays (MPRAs) conducted in THP1-derived macrophages (Bond et al., 2024). Both variants were found within a cCRE present in untreated and LPS+IFNG-treated iTF-Microglia and mapped to *BIN1* via DegCre, ABC, and brain microglia link. **rs10792832 (AD)** was found in a cCRE of untreated and LPS+IFNG-treated iTF-Microglia and mapped to *PICALM* via DegCre and brain microglia links. This variant has been associated with altered *PICALM* expression, disrupted PU.1 binding, lipid droplet accumulation, and impaired microglia phagocytosis (Kozlova et al., 2024). **rs6430538 (PD)** is an eQTL for *TMEM163* in microglia and AD brain tissues (Qiu et al., 2024; Young et al., 2021) which mapped via DegCre to *TMEM163* in untreated iTF-Microglia. Notably, it was also mapped to *TMEM163* via brain microglia link from AD samples.

To determine if variants might disrupt TF binding, we used motifbreakR (**Figure 5E; see also Table S7**). Predicted TFs included EGR1, a TF known to mark inflammatory enhancers in macrophages (Trizzino et al., 2021), and IRF1, which functions antagonistically with EGR1 during macrophage polarization (Chu et al., 2021). We also identified NF-κB and STAT TFs, which have been implicated in microglia-driven neurodegeneration (Siddiqui et al., 2024; Yun et al., 2021). RXRs, which promote amyloid beta (Aβ) clearance by enhancing microglia phagocytosis (Cramer et al., 2012; Yamanaka et al., 2012), were also identified. Several of these TFs were differentially expressed across conditions. For example, *EGR1*, *RXRA* and *MAFB* were downregulated upon LPS+IFNG stimulation, while *IRF1*, *NFKB1* and *SPI1* were upregulated (**Figure 5F; see also Table S1**).

We also identified variant-TF associations using ENCODE and BrainTF ChIP-seq datasets (**Table S6**), confirming known interactions such as PU.1 binding to PICALM-rs10792832 (Kozlova et al., 2024).

## Discussion

As KOLF2.1J iPSCs are becoming a standard line for modeling ADRD-associated variants, we addressed two key questions: (1) Do KOLF2.1J iTF-Microglia resemble other iPSC-derived microglia and primary human microglia? (2) Are they suitable for linking disease-risk variants to microglial TRNs? Briefly, the answer to both questions is “yes,” as discussed below.

We found that doxycycline-inducible iTF cassettes accelerated microglial differentiation, halving the time compared to growth factor-only protocols, including commercial kits.

Transcriptomic profiling showed that KOLF2.1J iTF-Microglia closely resemble other iPSC-derived microglia, especially WTC11 iTF-Microglia (Dräger et al., 2022), suggesting that differentiation protocol, rather than genetic background, primarily shapes transcriptional identity. In contrast, growth factor-only differentiated microglia (Abud et al., 2017) were more transcriptionally similar to primary fetal and adult microglia, indicating a more differentiated state. Nonetheless, both WTC11 and KOLF2.1J iTF-Microglia were more similar to primary microglia than to monocytes or induced hematopoietic progenitor cells (iHPCs), reinforcing their microglial identity. KOLF2.1J iTF-Microglia were also enriched for marker genes of microglia in human brain. Upon LPS+IFNG treatment, iTF-Microglia downregulated homeostatic genes and upregulated pathways associated with brain aging and neurodegeneration, such as interferon signaling, complement regulation, and antigen presentation (Hammond et al., 2019; Kann et al., 2022), highlighting their value for studying inflammatory microglial states.

Most cell-type-specific cCREs were located in distal enhancers, while shared cCREs across iTF-iPSCs, untreated, and LPS+IFNG-treated iTF-Microglia were enriched at promoter-proximal regions, underscoring the role of distal enhancers in cell-type and state-specific transcriptional regulation (Bulger and Groudine, 2011). iTF-iPSC cCREs were enriched for pluripotency pathways, while iTF-Microglia-specific cCREs were associated with innate immunity. Both untreated and LPS+IFNG-treated iTF-Microglia showed significantly greater overlap with brain microglia cCREs (Anderson et al., 2023) than iTF-iPSCs, highlighting their regulatory similarity to primary microglia.

LPS+IFNG stimulation led to chromatin opening at several cCREs without chromatin closure. H3K27ac changes included both gains and losses, such as reducing acetylation at the *TREM2* promoter that correlated with transcriptional repression, consistent with its known downregulation in inflammatory contexts (Liu et al., 2020). This repression may involve histone deacetylases (HDACs), which have been implicated in microglial activation (Kuboyama et al., 2017; Meleady et al., 2023). Notably, HDAC inhibition enhances Aβ phagocytosis and promotes disease-associated microglial states that may require TREM2 function (Datta et al., 2018; Haage et al., 2024).

TRNs, comprising cCREs, target genes and upstream TFs, were enriched for disease-risk variants. Genes expressed in untreated iTF-Microglia were associated with AD risk variants, suggesting that susceptibility pathways are active under homeostatic conditions, and that their repression under inflammatory environments may contribute to AD pathogenesis and progression. This aligns with previous studies showing that inflammatory stimuli downregulates the expression of AD-associated genes in microglia (Pulido-Salgado et al., 2018; Snijders et al., 2023).

In contrast, genes expressed in LPS+IFNG-treated iTF-Microglia were enriched for MS, IBD, and SLE variants, consistent with autoimmune risk loci being active in inflammatory conditions (Harroud and Hafler, 2023). Interestingly, these genes were also enriched for AD variants from the Jansen et al. GWAS (Jansen et al., 2019), potentially capturing unique loci found in this dataset, such as the *ADAMTS4*, *HESK1*, and *CNTNAP2* loci (Andrews et al., 2023).

iTF-iPSCs-specific cCREs were enriched for neuropsychiatric risk loci, suggesting developmental contributions to these disorders (Forrest et al., 2017). In contrast, untreated and LPS+IFNG-treated iTF-Microglia cCREs were enriched for AD and MS variants, consistent with findings from primary and in vitro microglia (Anderson et al., 2023; Novikova et al., 2021; Xiong et al., 2023). Mapping neurodegenerative risk loci to KOLF2.1J TRNs linked both known and novel variant-target gene associations, including *BIN1*, *TREM2* and *MS4A4A* (AD); *LRRK2* (PD); *HLA-DRB1* (MS); and *TBK1* (ALS/FTD) (Andrews et al., 2023; Harding et al., 2021; Kia et al., 2021; Yates et al., 2022)

We also identified TFs likely mediating disease risk. Several ETS family members (*SPI1*, ETVs, ELFs) were associated with AD variant enrichment (Kosoy et al., 2022), with *SPI1* being a known AD risk gene (Huang et al., 2017). MAFB and IKZF1 were also linked to AD (Anderson et al., 2023; Zhao et al., 2024), while NF-kB, IRF, and STAT TFs have been implicated in various neurodegenerative diseases (Hoenen et al., 2016; Sun et al., 2022; Yan et al., 2018). However, validation of variant-TF interactions in microglia remains limited, partly due to the scarcity of TF-specific ChIP-seq data (Kozlova et al., 2024).

This study has several limitations. First, analyses were limited to Day 18 of differentiation. Earlier time points, as explored with the WTC11 iTF line (Dräger et al., 2022), could help pinpoint when microglial TRNs are established, and identify optimal windows for high-throughput assays such as MPRAs or CRISPR screens. This is particularly relevant given the low transduction and transfection rate of differentiated microglia, which makes it challenging to apply these tools.

Performing these assays at the iPSC stage, when transduction rates are higher, and then differentiating into microglia may help circumvent this issue (Dräger et al., 2022). Notably, our preliminary data (not shown) indicate that CRISPRi targeting of promoters and cCREs, including a *BIN1*-linked microglial enhancer (Nott et al., 2019), is feasible as early as six days into iTF differentiation, when cells already express microglial markers.

Second, TRN inferences were based on computational predictions, which can introduce biases. Experimental validation, using tools such as MPRAs, CRISPR perturbations, or ChIP-seq, remains critical for refining these networks. Encouragingly, we successfully performed CETCh-seq in KOLF2.1J cells (not shown), enabling genome-wide TF binding profiling without the need for specific antibodies (Savic et al., 2015).

Finally, while molecular profiling was extensive, functional validation of iTF-Microglia was limited. Preliminary assays in KOLF2.1J iTF-Microglia confirmed phagocytosis of FITC-labeled beads, GPR34 protein expression (homeostatic marker), and CD38 protein upregulation (inflammatory marker) following LPS+IFNG treatment. However, these functional data were not included and should be further explored in future studies.

Overall, KOLF2.1J iTF-Microglia recapitulate TRNs of primary and in vitro microglia, providing a rapid, scalable, and cost-effective platform for genome-wide functional genomics. KOLF2.1J iTF-Microglia offer a valuable platform to study cell-autonomous effects of disease-risk variants on microglial gene regulation and function.

## Methods

### KOLF2.1J iTF-iPSC culture

KOLF2.1J CLYBL_6-TF-iMG_KI1/KI2 iPSCs (JIPSC002072), hereafter referred to as iTF-iPSCs, were obtained from The Jackson Laboratory. Cells were maintained at 37°C with 5% CO_2_ in mTeSR Plus medium (StemCell Technologies) on dishes coated with Growth Factor Reduced (GFR), LDEV-Free Matrigel (Gibco). Medium was changed every other day. Cells were passaged at 70–80% confluency using ReLeSR (StemCell Technologies), resuspended in room-temperature mTeSR Plus, and transferred to freshly coated dishes.

### KOLF2.1J iTF-Microglia differentiation

Differentiation was performed following Dräger et al. protocol (Dräger et al., 2022) with minor modifications. iTF-iPSCs were washed with D-PBS, dissociated with Accutase (StemCell Technologies), and centrifuged at 300×g for 5 minutes. Cell pellets were resuspended in Essential 8 Basal Medium (Gibco) supplemented with 10 μM Y-27632 ROCK Inhibitor and 2 μg/mL doxycycline (StemCell Technologies), then seeded onto double-coated plates with Poly-D-Lysine-precoated Bio plates (Corning) and GFR Matrigel.

On day 2, cells were switched to differentiation medium (Advanced DMEM/F12 (Gibco) supplemented with 1× GlutaMAX (Gibco), 2 μg/mL doxycycline, 100 ng/mL human IL-34 and 10 ng/mL human GM-CSF (PeproTech)). On day 4, the medium was replaced with iTF-Microglia medium (Advanced DMEM/F12 supplemented with 1× GlutaMAX, 2 μg/mL doxycycline, 100 ng/mL human IL-34, 10 ng/mL human GM-CSF, 50 ng/mL human M-CSF and 50 ng/mL human TGF-β1 (PeproTech). On day 8, fresh iTF-Microglia medium was added. Cells were maintained for 9 more days with full medium changes every 2–3 days.

### Lipopolysaccharide (LPS) and interferon-gamma (IFNG) treatment

iTF-iPSCs and iTF-Microglia on day 17 of differentiation were either untreated or stimulated with 100 ng/mL LPS (Invitrogen) and 20 ng/mL IFNG (Gibco) in fresh mTeSR Plus (for iTF-iPSCs) or iTF-Microglia medium (for iTF-Microglia) for 24 hours before harvesting.

### RNA-seq and data analysis

Total RNA was extracted from untreated and LPS+IFNG-treated iTF-iPSCs and iTF-Microglia (three replicates per condition) using the Total RNA Purification Plus Kit (Norgen). RNA concentration was measured with the Qubit RNA HS Assay Kit (ThermoFisher). Library preparation and paired-end 150 bp (PE150) sequencing were performed by Novogene on an Illumina NovaSeq X Plus platform.

Raw reads were quality-checked with FastQC before and after trimming. Trim Galore [paired-end mode, --illumina flag] was used to remove adapter sequences and low-quality bases. Trimmed reads were quantified using Salmon [--validateMappings, --seqBias, --gcBias] (Patro et al., 2017). Reads were mapped to the GENCODE v46 human transcriptome (hg38) using a k-mer size of 31.

Transcript-level quantifications were imported into R with tximeta (Love et al., 2020), and summarized to gene-level using summarizeToGene(). Differential expression analysis was performed with DESeq2 (Love et al., 2014) after pre-filtering genes with <10 counts in at least three samples. Differentially expressed genes (DEGs) were identified using the Wald test, with contrasts defined between iTF-iPSCs and iTF-Microglia in untreated and LPS+IFNG-stimulated conditions. Log2 fold changes (LFC) were shrunk using the ashr method (Stephens, 2016). Functional enrichment analysis was conducted with clusterProfiler (Yu et al., 2012) using Reactome pathway annotations (Yu and He, 2016).

### RNA-seq meta-analysis

Public RNA-seq datasets were obtained from the Gene Expression Omnibus (GEO) to analyze gene expression in iPSCs, iPSC-derived microglia, primary human microglia, and blood monocytes. Raw data from Abud et al. (GSE89189) (Abud et al., 2017) and Dräger et al. (GSE178317) (Dräger et al., 2022) were downloaded using the SRA Toolkit (v3.0.0) and converted to FASTQ with fasterq-dump, followed by gzip compression.

FASTQ files from Abud et al. were processed using the same pipeline as above. The Dräger et al. dataset, generated with the QuantSeq 3′ mRNA-Seq Library Prep Kit for Illumina (FWD) (Lexogen), was processed with a custom pipeline. Reads were trimmed with BBDuk to remove adapters and poly-A tail, quality checked with FastQC, and quantified using Salmon [-- validateMappings, --noLengthCorrection] and mapped to the GENCODE v46 human transcriptome (hg38) with a k-mer size of 27.

Gene-level counts were analyzed using DESeq2, applying size factor normalization to adjust for library size. Genes with ≥1 normalized count in ≥3 samples were retained. Variance-stabilizing transformation (VST) was applied to stabilize variance across expression levels. Marker genes for iPSCs (Dräger et al., 2022), microglia and blood monocytes (Abud et al., 2017) were used to assess cellular identity via hierarchical clustering.

To assess the transcriptional signatures of KOLF2.1J iTF cells in the context of brain cell types and microglial states, we used snRNAseq datasets from postmortem AD and control brains (Green et al., 2024; Mathys et al., 2024). Marker gene sets from the supplemental tables were formatted as GMT files for gene set enrichment analysis (GSEA) using fgsea (Korotkevich et al., 2016).

### ATAC-seq library preparation

ATAC-seq was performed on untreated iTF-iPSCs, and untreated and LPS+IFNG-treated iTF-Microglia (two replicates per condition) following a published protocol (Rogers et al., 2024). Briefly, 100,000 cells were harvested by mechanical dissociation, washed with 50 μL cold 1X PBS, and centrifuged at 300×g for 5 minutes. The pellet was resuspended in 50 μL cold lysis buffer (10 mM Tris-HCl pH 7.5, 10 mM NaCl, 3 mM MgCl_2_, 0.1% NP-40, 0.1% Tween-20, 0.01% Digitonin, and nuclease-free H_2_O) and incubated for 3 minutes on ice. After lysis, 1 mL of wash buffer (10 mM Tris-HCl pH 7.5, 10 mM NaCl, 3 mM MgCl_2_, 0.1% Tween-20, and nuclease-free H₂O) was added, mixed by gentle inversion, and centrifuged at 500×g for 10 minutes at 4°C. The nuclei pellet was resuspended in 50 μL transposition mix (Tagment DNA TDE1 Enzyme and Buffer, Illumina) and incubated at 37°C for 30 minutes with shaking (1000 rpm). DNA was purified using the Qiagen MinElute Reaction Cleanup Kit and eluted in 11 μL.

Libraries were generated using Nextera XT Index primers (Illumina) and Q5 Hot Start Master Mix (NEB), followed by PCR amplification (30 sec at 98°C; [10 sec at 98°C, 30 sec at 63°C, 1 min at 72°C] ×10 cycles). Libraries were quantified using the Qubit dsDNA HS Assay Kit and quality-checked with an Agilent BioAnalyzer High Sensitivity DNA chip. Sequencing was performed on an Illumina NovaSeq X Plus (PE150), targeting ∼200 million reads per library.

### H3K27ac ChIP-seq library preparation

ChIP-seq for H3K27ac was performed on chromatin from untreated iTF-iPSCs, and untreated and LPS+IFNG-treated iTF-Microglia (two replicates per condition), following protocols from our lab and the ENCODE Consortium. Chromatin was immunoprecipitated with an anti-H3K27ac antibody (ActiveMotif). Libraries were prepared by end-repair, adapter ligation, and PCR amplification (30 sec at 98°C; [10 sec at 98°C, 30 sec at 65°C, 30 sec at 72°C] ×15 cycles; 5 min at 72°C). Libraries were quantified with the Qubit dsDNA HS Assay Kit and analyzed with the Standard Sensitivity NGS Fragment Analysis Kit (Advanced Analytical) on an Agilent Fragment Analyzer 5200. Sequencing was performed on an Illumina NovaSeq X Plus (PE150), targeting ∼30 million reads per library.

### ATAC- and ChIP-seq processing and peak calling

ATAC-seq data were processed with the ENCODE ATAC-seq pipeline (v1.7.0). Reads were trimmed with Cutadapt, aligned to the hg38 reference genome using Bowtie2, and filtered to remove duplicates and mitochondrial reads using Picard and Samtools. High-quality, properly paired reads were retained. Peaks were called with MACS2 [--shift −75 --extsize 150], filtered for ENCODE blacklist regions, and reproducible peaks identified using IDR and naive overlap.

ChIP-seq reads were pre-processed with clumpify (BBMap v38.42) to remove optical duplicates [dedupe=t optical=t dupedist=12000] and processed with the ENCODE chip-seq-pipeline2 with histone-specific settings [-type histone]. Reads were trimmed with Trimmomatic, aligned to the hg38 reference genome using Bowtie2, and PCR duplicates removed with Picard. Peaks were called with MACS2, and reproducible peaks identified via naive overlap.

### Hi-C Chromosome Conformation Capture Assays

Hi-C libraries were generated using the Arima-HiC+ Genome-Wide Hi-C kit (Arima Genomics) following the manufacturer’s protocol. Chromatin from untreated iTF-iPSCs, untreated iTF-Microglia, and LPS+IFNG-treated iTF-Microglia (two replicates per condition) was crosslinked, digested with restriction enzymes, and proximity-ligated to capture 3D chromatin interactions. Ligated DNA was purified, fragmented, and prepared for sequencing. A pooled 6-plex library was sequenced on an Illumina NovaSeq X Plus 25B lane (PE150), generating 3.3 billion paired-end reads (∼550 million reads per library).

Hi-C data were processed using the ENCODE Hi-C pipeline (v1.15.1) following ENCODE standards. Sequencing reads were aligned to the hg38 genome using BWA-MEM. Low-quality, secondary, and duplicate reads were removed, and valid interaction pairs were retained to construct contact matrices at multiple resolutions. A combined .hic file was generated by merging all samples across cell types (iTF-iPSCs and iTF-Microglia) and conditions (untreated and LPS+IFNG-treated).

Chromatin loops were identified using HiCCUPS, as implemented in the ENCODE Hi-C pipeline. Loops were called from the combined .hic file. Final loop coordinates were extracted from the 30.bedpe.gz output file.

### Nomination of Transcriptional Regulatory Networks (TRNs)

#### Identification of Candidate Cis-Regulatory Elements (cCREs)

cCREs were nominated by overlapping chromatin accessibility (ATAC-seq) and enhancer-associated histone modifications (H3K27ac ChIP-seq). Optimal reproducible peaks from two replicates per dataset were filtered (q-value ≥ 2) and overlapped using findOverlaps from the GenomicRanges package.

To standardize genomic coordinates across cell types and states, we generated a consensus cCRE set by merging cCREs identified in each condition. Cell-type specific cCREs were then mapped to this consensus set, retaining the consensus coordinates.

Functional enrichment of cCREs was performed using rGREAT (Gu and Hübschmann, 2023) with the C2:CP:Reactome pathway collection from the Human Molecular Signatures Database (MSigDB) (Milacic et al., 2024).

Differential accessibility and H3K27ac marking were assessed using the csaw package (Lun and Smyth, 2016). ATAC-seq and ChIP-seq reads were quantified within cCREs using regionCounts, normalized using trimmed mean of M-values (TMM) with background bins generated by counting reads in 10 kb non-overlapping windows (windowCounts) across the genome. A generalized linear model (GLM) was applied to assess differential accessibility and H3K27ac marking between conditions.

#### Identification of TFs

To predict TFs driving differential chromatin accessibility during microglial differentiation (iTF-iPSCs vs. iTF-Microglia) and H3K27ac changes upon inflammatory stimulation (LPS+IFNG-treated iTF-Microglia vs. iTF-Microglia), we applied the monaLisa package (Machlab et al., 2022). Specifically, we used the randomized lasso stability selection approach to identify TF motifs enriched in cCREs showing changes in accessibility or H3K27ac levels, based on results from csaw. Motif scanning was performed using JASPAR2024 position weight matrices (PWMs) via findMotifHits, with GC content and observed/expected CpG ratios included as covariates to account for sequence bias. TFs were considered significant if their selection probability was ≥ 0.8.

#### Identification of Target Genes

We used four complementary approaches to assign target genes to cCREs and TFs: (1) **Closest Gene Assignment:** Each cCRE was assigned to its nearest gene TSS (GENCODE v46, hg38) using nearest() and distanceToNearest(), restricted to genes with RNA-seq expression > 1 TPM. (2) **DegCre Analysis:** Using DegCre (Roberts et al., 2024), we correlated DEGs (from DESeq2) with cCREs showing differential accessibility during microglia differentiation (iTF-Microglia vs iTF-iPSCs) and H3K27ac changes after stimulation (iTF-Microglia-treated LPS+IFNG vs iTF-Microglia), from csaw analysis. Only associations with concordant LFC direction and assocAlpha ≤ 0.05 were retained. (3) **ABC Model:** We applied the ABC model (Fulco et al., 2019) to predict enhancer-gene links based on enhancer activity (ATAC and H3K27ac datasets) and Hi-C contact frequencies (combined .hic file). Predicted enhancer-gene associations were filtered according to published ABC score thresholds (Nasser et al., 2021), retaining only promoters with scores ≥ 0.1 and enhancers with scores ≥ 0.015. (4) **Brain microglia links:** We leveraged snMULTI-omics data from post-mortem brains of individuals with and without AD (Anderson et al., 2023). cCRE-to-gene links were inferred by correlating chromatin accessibility and gene expression within the same nucleus. To maintain relevance to our system, we retained only links involving brain microglia and overlapping our iTF-derived cCREs.

Functional enrichment analysis of identified target genes was conducted with clusterProfiler (Yu et al., 2012) using Reactome pathway annotations (Yu and He, 2016).

### Association of TRNs with candidate disease-risk variants

To link TRNs with disease-risk variants, we performed:

MAGMA (v1.10) analysis (De Leeuw et al., 2015). GWAS variants were mapped to gene bodies with a 2 kb upstream window from the TSS. To account for expression-level effects, we performed conditional gene expression analysis using log2(TPM+1) values, filtered for TPM > 1, winsorized at 50 TPM, and included as covariates.

sLDSC (v.1.0.1) analysis with the full baseline-LD v2.2 model (Bulik-Sullivan et al., 2015; Finucane et al., 2015) to test for enrichment of GWAS variants in ATAC and H3K27ac peaks, cCREs and TF-defined cCRE subsets (from monaLisa).

GWAS summary statistics used for MAGMA and sLDSC analyses are listed in **Table S8**. For AD GWAS summary statistics, the *APOE* and *HLA* regions were excluded.

To investigate the relevance to neurodegenerative disorders, we intersected TRNs with genetic variants linked to AD, ALS, LBD, MS, PD, frontotemporal dementia (FTD), Huntington’s disease (HD), multiple system atrophy (MSA), and progressive supranuclear palsy (PSP) from the NHGRI-EBI GWAS Catalog (Sollis et al., 2023). Given that GWAS-identified variants are often proxies in LD with causal variants, we curated 3,985 GWAS-identified variants and used TOP-LD (Huang et al., 2022) to query the TOPMed dataset (Taliun et al., 2021), selecting variants with minor allele frequency (MAF) > 0.01 and LD threshold (R² > 0.8). This resulted in a set of 62,300 LD-linked candidate variants, which we intersected with cCREs from iTF-iPSCs, untreated iTF-Microglia, and LPS+IFNG-treated iTF-Microglia.

Target genes were assigned to candidate variants using the closest active gene, DegCre, ABC predictions, and brain microglia links. Variant annotations included microglia eQLTs (Lopes et al., 2022; Young et al., 2021) and MPRAs in THP1-derived macrophages under resting and LPS+IFNG-stimulated conditions (Bond et al., 2024).

To identify TFs whose binding may be disrupted by variants, we overlapped variant locations with: (1) **ENCODE ChIP-seq:** TF binding data from 1,151 TFs across 188 biosamples, including cell lines, in vitro differentiated cells, organoids, primary cells, and tissues (ENCODE Project Consortium et al., 2020; Luo et al., 2020). (2) **BrainTF:** ChIP-seq maps for >100 TFs in post-mortem brain tissues and sorted nuclei, including NeuN+ (neurons), Olig+ (oligodendrocytes), and double-negative nuclei (primarily microglia and astrocytes) (Loupe et al., 2024). Peaks from double-negative nuclei were analyzed separately to infer microglia-enriched TF binding. (3) **motifbreakR:** TF motif disruption was predicted using the “ic” algorithm and PWMs from motifbreakR_motif, with a P-value cutoff of < 1×10⁻^4^ (Coetzee et al., 2015).

## References

Abdelalim, E.M., Emara, M.M., and Kolatkar, P.R. (2014). The SOX transcription factors as key players in pluripotent stem cells. Stem Cells Dev 23, 2687–2699. 10.1089/scd.2014.0297.

Abud, E.M., Ramirez, R.N., Martinez, E.S., Healy, L.M., Nguyen, C.H.H., Newman, S.A., Yeromin, A.V., Scarfone, V.M., Marsh, S.E., Fimbres, C., et al. (2017). iPSC-Derived Human Microglia-like Cells to Study Neurological Diseases. Neuron 94, 278–293.e9. 10.1016/j.neuron.2017.03.042.

Adams, L., Song, M.K., Yuen, S., Tanaka, Y., and Kim, Y.-S. (2024). A single-nuclei paired multiomic analysis of the human midbrain reveals age- and Parkinson’s disease-associated glial changes. Nat Aging 4, 364–378. 10.1038/s43587-024-00583-6.

Anderson, A.G., Rogers, B.B., Loupe, J.M., Rodriguez-Nunez, I., Roberts, S.C., White, L.M., Brazell, J.N., Bunney, W.E., Bunney, B.G., Watson, S.J., et al. (2023). Single nucleus multiomics identifies ZEB1 and MAFB as candidate regulators of Alzheimer’s disease-specific cis-regulatory elements. Cell Genomics 3. 10.1016/j.xgen.2023.100263.

Andrews, S.J., Renton, A.E., Fulton-Howard, B., Podlesny-Drabiniok, A., Marcora, E., and Goate, A.M. (2023). The complex genetic architecture of Alzheimer’s disease: novel insights and future directions. eBioMedicine 90, 104511. 10.1016/j.ebiom.2023.104511.

Askarova, A., Yaa, R.M., Marzi, S.J., and Nott, A. (2024). Genetic risk for neurodegenerative conditions is linked to disease-specific microglial pathways. 10.1101/2024.08.29.610255.

Ballasch, I., García-García, E., Vila, C., Pérez-González, A., Sancho-Balsells, A., Fernández, J., Soto, D., Puigdellívol, M., Gasull, X., Alberch, J., et al. (2023). Ikzf1 as a novel regulator of microglial homeostasis in inflammation and neurodegeneration. Brain, Behavior, and Immunity 109, 144–161. 10.1016/j.bbi.2023.01.016.

Bond, M.L., Quiroga-Barber, I.Y., D’Costa, S., Wu, Y., Bell, J.L., McAfee, J.C., Kramer, N.E., Lee, S., Patrucco, M., Phanstiel, D.H., et al. (2024). Deciphering the functional impact of Alzheimer’s Disease-associated variants in resting and proinflammatory immune cells. medRxiv 2024.09.13.24313654. 10.1101/2024.09.13.24313654.

Brenner, D., Sieverding, K., Bruno, C., Lüningschrör, P., Buck, E., Mungwa, S., Fischer, L., Brockmann, S.J., Ulmer, J., Bliederhäuser, C., et al. (2019). Heterozygous Tbk1 loss has opposing effects in early and late stages of ALS in mice. J Exp Med 216, 267–278. 10.1084/jem.20180729.

Bulger, M., and Groudine, M. (2011). Functional and mechanistic diversity of distal transcription enhancers. Cell 144, 327–339. 10.1016/j.cell.2011.01.024.

Bulik-Sullivan, B.K., Loh, P.-R., Finucane, H.K., Ripke, S., Yang, J., Schizophrenia Working Group of the Psychiatric Genomics Consortium, Patterson, N., Daly, M.J., Price, A.L., and Neale, B.M. (2015). LD Score regression distinguishes confounding from polygenicity in genome-wide association studies. Nat Genet 47, 291–295. 10.1038/ng.3211.

Buttgereit, A., Lelios, I., Yu, X., Vrohlings, M., Krakoski, N.R., Gautier, E.L., Nishinakamura, R., Becher, B., and Greter, M. (2016). Sall1 is a transcriptional regulator defining microglia identity and function. Nat Immunol 17, 1397–1406. 10.1038/ni.3585.

Carrasquillo, M.M., Allen, M., Burgess, J.D., Wang, X., Strickland, S.L., Aryal, S., Siuda, J., Kachadoorian, M.L., Medway, C., Younkin, C.S., et al. (2017). A candidate regulatory variant at the TREM gene cluster associates with decreased Alzheimer’s disease risk and increased TREML1 and TREM2 brain gene expression. Alzheimers Dement 13, 663–673. 10.1016/j.jalz.2016.10.005.

Carter, S.L., Müller, M., Manders, P.M., and Campbell, I.L. (2007). Induction of the genes for Cxcl9 and Cxcl10 is dependent on IFN-gamma but shows differential cellular expression in experimental autoimmune encephalomyelitis and by astrocytes and microglia in vitro. Glia 55, 1728–1739. 10.1002/glia.20587.

Chen, S.-W., Hung, Y.-S., Fuh, J.-L., Chen, N.-J., Chu, Y.-S., Chen, S.-C., Fann, M.-J., and Wong, Y.-H. (2021). Efficient conversion of human induced pluripotent stem cells into microglia by defined transcription factors. Stem Cell Reports 16, 1363–1380. 10.1016/j.stemcr.2021.03.010.

Chu, Y., Li, J., Jia, P., Cui, J., Zhang, R., Kang, X., Lv, M., and Zhang, S. (2021). Irf1- and Egr1-activated transcription plays a key role in macrophage polarization: A multiomics sequencing study with partial validation. International Immunopharmacology 99, 108072. 10.1016/j.intimp.2021.108072.

Cirulli, E.T., Lasseigne, B.N., Petrovski, S., Sapp, P.C., Dion, P.A., Leblond, C.S., Couthouis, J., Lu, Y.-F., Wang, Q., Krueger, B.J., et al. (2015). Exome sequencing in amyotrophic lateral sclerosis identifies risk genes and pathways. Science 347, 1436–1441. 10.1126/science.aaa3650.

Coetzee, S.G., Coetzee, G.A., and Hazelett, D.J. (2015). motifbreakR: an R/Bioconductor package for predicting variant effects at transcription factor binding sites. Bioinformatics 31, 3847–3849. 10.1093/bioinformatics/btv470.

Cramer, P.E., Cirrito, J.R., Wesson, D.W., Lee, C.Y.D., Karlo, J.C., Zinn, A.E., Casali, B.T., Restivo, J.L., Goebel, W.D., James, M.J., et al. (2012). ApoE-directed therapeutics rapidly clear β-amyloid and reverse deficits in AD mouse models. Science 335, 1503–1506. 10.1126/science.1217697.

Datta, M., Staszewski, O., Raschi, E., Frosch, M., Hagemeyer, N., Tay, T.L., Blank, T., Kreutzfeldt, M., Merkler, D., Ziegler-Waldkirch, S., et al. (2018). Histone Deacetylases 1 and 2 Regulate Microglia Function during Development, Homeostasis, and Neurodegeneration in a Context-Dependent Manner. Immunity 48, 514–529.e6. 10.1016/j.immuni.2018.02.016.

De Leeuw, C.A., Mooij, J.M., Heskes, T., and Posthuma, D. (2015). MAGMA: Generalized Gene-Set Analysis of GWAS Data. PLoS Comput Biol 11, e1004219. 10.1371/journal.pcbi.1004219.

Dobner, J., Nguyen, T., Dunkel, A., Prigione, A., Krutmann, J., and Rossi, A. (2024a). Mitochondrial DNA integrity and metabolome profile are preserved in the human induced pluripotent stem cell reference line KOLF2.1J. Stem Cell Reports 19, 343–350. 10.1016/j.stemcr.2024.01.009.

Dobner, J., Diecke, S., Krutmann, J., Prigione, A., and Rossi, A. (2024b). Reassessment of marker genes in human induced pluripotent stem cells for enhanced quality control. Nat Commun 15, 8547. 10.1038/s41467-024-52922-1.

Dou, D., Holzbaur, E.L.F., and Boecker, C.A. (2025). Protocol for live imaging of axonal transport in iPSC-derived iNeurons. STAR Protoc 6, 103556. 10.1016/j.xpro.2024.103556.

Douvaras, P., Sun, B., Wang, M., Kruglikov, I., Lallos, G., Zimmer, M., Terrenoire, C., Zhang, B., Gandy, S., Schadt, E., et al. (2017). Directed Differentiation of Human Pluripotent Stem Cells to Microglia. Stem Cell Reports 8, 1516–1524. 10.1016/j.stemcr.2017.04.023.

Dräger, N.M., Sattler, S.M., Huang, C.T.-L., Teter, O.M., Leng, K., Hashemi, S.H., Hong, J., Aviles, G., Clelland, C.D., Zhan, L., et al. (2022). A CRISPRi/a platform in human iPSC-derived microglia uncovers regulators of disease states. Nat Neurosci 25, 1149–1162. 10.1038/s41593-022-01131-4.

Drummond, E., and Wisniewski, T. (2017). Alzheimer’s disease: experimental models and reality. Acta Neuropathol 133, 155–175. 10.1007/s00401-016-1662-x.

ENCODE Project Consortium, Moore, J.E., Purcaro, M.J., Pratt, H.E., Epstein, C.B., Shoresh, N., Adrian, J., Kawli, T., Davis, C.A., Dobin, A., et al. (2020). Expanded encyclopaedias of DNA elements in the human and mouse genomes. Nature 583, 699–710. 10.1038/s41586-020-2493-4.

Fang, K., Pishva, E., Piers, T., and Scholpp, S. (2025). Amyloid-β can activate JNK signalling via WNT5A-ROR2 to reduce synapse formation in Alzheimer’s disease. J Cell Sci 138, JCS263526. 10.1242/jcs.263526.

Field, A., and Adelman, K. (2020). Evaluating Enhancer Function and Transcription. Annu. Rev. Biochem. 89, 213–234. 10.1146/annurev-biochem-011420-095916.

Finucane, H.K., Bulik-Sullivan, B., Gusev, A., Trynka, G., Reshef, Y., Loh, P.-R., Anttila, V., Xu, H., Zang, C., Farh, K., et al. (2015). Partitioning heritability by functional annotation using genome-wide association summary statistics. Nat Genet 47, 1228–1235. 10.1038/ng.3404.

Fixsen, B.R., Han, C.Z., Zhou, Y., Spann, N.J., Saisan, P., Shen, Z., Balak, C., Sakai, M., Cobo, I., Holtman, I.R., et al. (2023). SALL1 enforces microglia-specific DNA binding and function of SMADs to establish microglia identity. Nat Immunol 24, 1188–1199. 10.1038/s41590-023-01528-8.

Forrest, M.P., Zhang, H., Moy, W., McGowan, H., Leites, C., Dionisio, L.E., Xu, Z., Shi, J., Sanders, A.R., Greenleaf, W.J., et al. (2017). Open Chromatin Profiling in hiPSC-Derived Neurons Prioritizes Functional Noncoding Psychiatric Risk Variants and Highlights Neurodevelopmental Loci. Cell Stem Cell 21, 305–318.e8. 10.1016/j.stem.2017.07.008.

Fulco, C.P., Nasser, J., Jones, T.R., Munson, G., Bergman, D.T., Subramanian, V., Grossman, S.R., Anyoha, R., Doughty, B.R., Patwardhan, T.A., et al. (2019). Activity-by-contact model of enhancer-promoter regulation from thousands of CRISPR perturbations. Nat Genet 51, 1664–1669. 10.1038/s41588-019-0538-0.

Gamache, J., Gingerich, D., Shwab, E.K., Barrera, J., Garrett, M.E., Hume, C., Crawford, G.E., Ashley-Koch, A.E., and Chiba-Falek, O. (2023). Integrative single-nucleus multi-omics analysis prioritizes candidate cis and trans regulatory networks and their target genes in Alzheimer’s disease brains. Cell Biosci 13, 185. 10.1186/s13578-023-01120-5.

Gao, C., Jiang, J., Tan, Y., and Chen, S. (2023). Microglia in neurodegenerative diseases: mechanism and potential therapeutic targets. Sig Transduct Target Ther 8, 359. 10.1038/s41392-023-01588-0.

Gao, T., Jernigan, J., Raza, S.A., Dammer, E.B., Xiao, H., Seyfried, N.T., Levey, A.I., and Rangaraju, S. (2019). Transcriptional regulation of homeostatic and disease-associated-microglial genes by IRF1, LXRβ, and CEBPα. Glia 67, 1958–1975. 10.1002/glia.23678.

Gerrits, E., Heng, Y., Boddeke, E.W.G.M., and Eggen, B.J.L. (2020). Transcriptional profiling of microglia; current state of the art and future perspectives. Glia 68, 740–755. 10.1002/glia.23767.

Gosselin, D., Skola, D., Coufal, N.G., Holtman, I.R., Schlachetzki, J.C.M., Sajti, E., Jaeger, B.N., O’Connor, C., Fitzpatrick, C., Pasillas, M.P., et al. (2017). An environment-dependent transcriptional network specifies human microglia identity. Science 356, eaal3222. 10.1126/science.aal3222.

Gracia-Diaz, C., Perdomo, J.E., Khan, M.E., Roule, T., Disanza, B.L., Cajka, G.G., Lei, S., Gagne, A.L., Maguire, J.A., Shalem, O., et al. (2024). KOLF2.1J iPSCs carry CNVs associated with neurodevelopmental disorders. Cell Stem Cell 31, 288–289. 10.1016/j.stem.2024.02.007.

Green, G.S., Fujita, M., Yang, H.-S., Taga, M., Cain, A., McCabe, C., Comandante-Lou, N., White, C.C., Schmidtner, A.K., Zeng, L., et al. (2024). Cellular communities reveal trajectories of brain ageing and Alzheimer’s disease. Nature 633, 634–645. 10.1038/s41586-024-07871-6.

Gu, Z., and Hübschmann, D. (2023). rGREAT: an R/bioconductor package for functional enrichment on genomic regions. Bioinformatics 39, btac745. 10.1093/bioinformatics/btac745.

Guttikonda, S.R., Sikkema, L., Tchieu, J., Saurat, N., Walsh, R.M., Harschnitz, O., Ciceri, G., Sneeboer, M., Mazutis, L., Setty, M., et al. (2021). Fully defined human pluripotent stem cell-derived microglia and tri-culture system model C3 production in Alzheimer’s disease. Nat Neurosci 24, 343–354. 10.1038/s41593-020-00796-z.

Haage, V., Tuddenham, J.F., Bautista, A., White, C.C., Garcia, F., Patel, R., Comandante-Lou, N., Marshe, V., Soni, R.K., Sims, P.A., et al. (2024). HDAC Inhibitors recapitulate Human Disease-Associated Microglia Signatures in vitro. bioRxiv 2024.10.11.617544. 10.1101/2024.10.11.617544.

Hammond, T.R., Marsh, S.E., and Stevens, B. (2019). Immune Signaling in Neurodegeneration. Immunity 50, 955–974. 10.1016/j.immuni.2019.03.016.

Harding, O., Evans, C.S., Ye, J., Cheung, J., Maniatis, T., and Holzbaur, E.L.F. (2021). ALS- and FTD-associated missense mutations in TBK1 differentially disrupt mitophagy. Proc Natl Acad Sci U S A 118, e2025053118. 10.1073/pnas.2025053118.

Harroud, A., and Hafler, D.A. (2023). Common genetic factors among autoimmune diseases. Science 380, 485–490. 10.1126/science.adg2992.

Herrera-Uribe, J., Liu, H., Byrne, K.A., Bond, Z.F., Loving, C.L., and Tuggle, C.K. (2020). Changes in H3K27ac at Gene Regulatory Regions in Porcine Alveolar Macrophages Following LPS or PolyIC Exposure. Front Genet 11, 817. 10.3389/fgene.2020.00817.

Hoenen, C., Gustin, A., Birck, C., Kirchmeyer, M., Beaume, N., Felten, P., Grandbarbe, L., Heuschling, P., and Heurtaux, T. (2016). Alpha-Synuclein Proteins Promote Pro-Inflammatory Cascades in Microglia: Stronger Effects of the A53T Mutant. PLoS One 11, e0162717. 10.1371/journal.pone.0162717.

Huang, K.-L., Marcora, E., Pimenova, A.A., Di Narzo, A.F., Kapoor, M., Jin, S.C., Harari, O., Bertelsen, S., Fairfax, B.P., Czajkowski, J., et al. (2017). A common haplotype lowers PU.1 expression in myeloid cells and delays onset of Alzheimer’s disease. Nat Neurosci 20, 1052–1061. 10.1038/nn.4587.

Huang, L., Rosen, J.D., Sun, Q., Chen, J., Wheeler, M.M., Zhou, Y., Min, Y.-I., Kooperberg, C., Conomos, M.P., Stilp, A.M., et al. (2022). TOP-LD: A tool to explore linkage disequilibrium with TOPMed whole-genome sequence data. The American Journal of Human Genetics 109, 1175–1181. 10.1016/j.ajhg.2022.04.006.

Huang, M., Malovic, E., Ealy, A., Jin, H., Anantharam, V., Kanthasamy, A., and Kanthasamy, A.G. (2023). Microglial immune regulation by epigenetic reprogramming through histone H3K27 acetylation in neuroinflammation. Front Immunol 14, 1052925. 10.3389/fimmu.2023.1052925.

Huang, P., Zhu, J., Liu, Y., Liu, G., Zhang, R., Li, D., Pei, D., and Zhu, P. (2021). Identification of New Transcription Factors that Can Promote Pluripotent Reprogramming. Stem Cell Rev and Rep 17, 2223–2234. 10.1007/s12015-021-10220-z.

Jaffe, A.B., and Hall, A. (2005). RHO GTPASES: Biochemistry and Biology. Annu. Rev. Cell Dev. Biol. 21, 247–269. 10.1146/annurev.cellbio.21.020604.150721.

Jansen, I.E., Savage, J.E., Watanabe, K., Bryois, J., Williams, D.M., Steinberg, S., Sealock, J., Karlsson, I.K., Hägg, S., Athanasiu, L., et al. (2019). Genome-wide meta-analysis identifies new loci and functional pathways influencing Alzheimer’s disease risk. Nat Genet 51, 404–413. 10.1038/s41588-018-0311-9.

Kaikkonen, M.U., Spann, N.J., Heinz, S., Romanoski, C.E., Allison, K.A., Stender, J.D., Chun, H.B., Tough, D.F., Prinjha, R.K., Benner, C., et al. (2013). Remodeling of the enhancer landscape during macrophage activation is coupled to enhancer transcription. Mol Cell 51, 310–325. 10.1016/j.molcel.2013.07.010.

Kann, O., Almouhanna, F., and Chausse, B. (2022). Interferon γ: a master cytokine in microglia-mediated neural network dysfunction and neurodegeneration. Trends Neurosci 45, 913–927. 10.1016/j.tins.2022.10.007.

Kia, D.A., Zhang, D., Guelfi, S., Manzoni, C., Hubbard, L., Reynolds, R.H., Botía, J., Ryten, M., Ferrari, R., Lewis, P.A., et al. (2021). Identification of Candidate Parkinson Disease Genes by Integrating Genome-Wide Association Study, Expression, and Epigenetic Data Sets. JAMA Neurol 78, 464. 10.1001/jamaneurol.2020.5257.

Konttinen, H., Cabral-da-Silva, M.E.C., Ohtonen, S., Wojciechowski, S., Shakirzyanova, A., Caligola, S., Giugno, R., Ishchenko, Y., Hernández, D., Fazaludeen, M.F., et al. (2019). PSEN1ΔE9, APPswe, and APOE4 Confer Disparate Phenotypes in Human iPSC-Derived Microglia. Stem Cell Reports 13, 669–683. 10.1016/j.stemcr.2019.08.004.

Korotkevich, G., Sukhov, V., Budin, N., Shpak, B., Artyomov, M.N., and Sergushichev, A. (2016). Fast gene set enrichment analysis. 10.1101/060012.

Kosoy, R., Fullard, J.F., Zeng, B., Bendl, J., Dong, P., Rahman, S., Kleopoulos, S.P., Shao, Z., Girdhar, K., Humphrey, J., et al. (2022). Genetics of the human microglia regulome refines Alzheimer’s disease risk loci. Nat Genet 54, 1145–1154. 10.1038/s41588-022-01149-1.

Kozlova, A., Zhang, S., Sudwarts, A., Zhang, H., Smirnou, S., Sun, X., Stephenson, K., Zhao, X., Jamison, B., Ponnusamy, M., et al. (2024). Alzheimer’s disease risk allele of PICALM causes detrimental lipid droplets in microglia. Res Sq rs.3.rs-4407146. 10.21203/rs.3.rs-4407146/v1.

Kracht, L., Borggrewe, M., Eskandar, S., Brouwer, N., Chuva De Sousa Lopes, S.M., Laman, J.D., Scherjon, S.A., Prins, J.R., Kooistra, S.M., and Eggen, B.J.L. (2020). Human fetal microglia acquire homeostatic immune-sensing properties early in development. Science 369, 530–537. 10.1126/science.aba5906.

Kuboyama, T., Wahane, S., Huang, Y., Zhou, X., Wong, J.K., Koemeter-Cox, A., Martini, M., Friedel, R.H., and Zou, H. (2017). HDAC3 inhibition ameliorates spinal cord injury by immunomodulation. Sci Rep 7, 8641. 10.1038/s41598-017-08535-4.

Langston, R.G., Beilina, A., Reed, X., Kaganovich, A., Singleton, A.B., Blauwendraat, C., Gibbs, J.R., and Cookson, M.R. (2022). Association of a common genetic variant with Parkinson’s disease is mediated by microglia. Sci. Transl. Med. 14, eabp8869. 10.1126/scitranslmed.abp8869.

Lee, A.J., Kim, C., Park, S., Joo, J., Choi, B., Yang, D., Jun, K., Eom, J., Lee, S.-J., Chung, S.J., et al. (2023). Characterization of altered molecular mechanisms in Parkinson’s disease through cell type-resolved multiomics analyses. Sci Adv 9, eabo2467. 10.1126/sciadv.abo2467.

Lemmens, M., Perner, J., Potgeter, L., Zogg, M., Thiruchelvam, S., Müller, M., Doll, T., Werner, A., Gilbart, Y., Couttet, P., et al. (2023). Identification of marker genes to monitor residual iPSCs in iPSC-derived products. Cytotherapy 25, 59–67. 10.1016/j.jcyt.2022.09.010.

Li, Y.E., Preissl, S., Miller, M., Johnson, N.D., Wang, Z., Jiao, H., Zhu, C., Wang, Z., Xie, Y., Poirion, O., et al. (2023). A comparative atlas of single-cell chromatin accessibility in the human brain. Science 382, eadf7044. 10.1126/science.adf7044.

Liu, W., Taso, O., Wang, R., Bayram, S., Graham, A.C., Garcia-Reitboeck, P., Mallach, A., Andrews, W.D., Piers, T.M., Botia, J.A., et al. (2020). Trem2 promotes anti-inflammatory responses in microglia and is suppressed under pro-inflammatory conditions. Hum Mol Genet 29, 3224–3248. 10.1093/hmg/ddaa209.

Liu, X., Wang, M., Jiang, T., He, J., Fu, X., and Xu, Y. (2019). IDO1 Maintains Pluripotency of Primed Human Embryonic Stem Cells by Promoting Glycolysis. Stem Cells 37, 1158–1165. 10.1002/stem.3044.

Lopes, K. de P., Snijders, G.J.L., Humphrey, J., Allan, A., Sneeboer, M.A.M., Navarro, E., Schilder, B.M., Vialle, R.A., Parks, M., Missall, R., et al. (2022). Genetic analysis of the human microglial transcriptome across brain regions, aging and disease pathologies. Nat Genet 54, 4–17. 10.1038/s41588-021-00976-y.

Loupe, J.M., Anderson, A.G., Rizzardi, L.F., Rodriguez-Nunez, I., Moyers, B., Trausch-Lowther, K., Jain, R., Bunney, W.E., Bunney, B.G., Cartagena, P., et al. (2024). Multiomic profiling of transcription factor binding and function in human brain. Nat Neurosci 10.1038/s41593-024-01658-8.

Love, M.I., Huber, W., and Anders, S. (2014). Moderated estimation of fold change and dispersion for RNA-seq data with DESeq2. Genome Biol 15, 550. 10.1186/s13059-014-0550-8.

Love, M.I., Soneson, C., Hickey, P.F., Johnson, L.K., Pierce, N.T., Shepherd, L., Morgan, M., and Patro, R. (2020). Tximeta: Reference sequence checksums for provenance identification in RNA-seq. PLoS Comput Biol 16, e1007664. 10.1371/journal.pcbi.1007664.

Lun, A.T.L., and Smyth, G.K. (2016). csaw: a Bioconductor package for differential binding analysis of ChIP-seq data using sliding windows. Nucleic Acids Res 44, e45. 10.1093/nar/gkv1191.

Luo, Y., Hitz, B.C., Gabdank, I., Hilton, J.A., Kagda, M.S., Lam, B., Myers, Z., Sud, P., Jou, J., Lin, K., et al. (2020). New developments on the Encyclopedia of DNA Elements (ENCODE) data portal. Nucleic Acids Res 48, D882–D889. 10.1093/nar/gkz1062.

Machlab, D., Burger, L., Soneson, C., Rijli, F.M., Schübeler, D., and Stadler, M.B. (2022). monaLisa: an R/Bioconductor package for identifying regulatory motifs. Bioinformatics 38, 2624–2625. 10.1093/bioinformatics/btac102.

Martinez-Sanchez, M., Skarnes, W., Jain, A., Vemula, S., Sun, L., Rockowitz, S., and Whitman, M.C. (2025). Chromosome 4 Duplication Associated with Strabismus Leads to Gene Expression Changes in iPSC-Derived Cortical Neurons. Genes (Basel) 16, 80. 10.3390/genes16010080.

Mathys, H., Boix, C.A., Akay, L.A., Xia, Z., Davila-Velderrain, J., Ng, A.P., Jiang, X., Abdelhady, G., Galani, K., Mantero, J., et al. (2024). Single-cell multiregion dissection of Alzheimer’s disease. Nature 632, 858–868. 10.1038/s41586-024-07606-7.

McLellan, A.D., Heiser, A., and Hart, D.N. (1999). Induction of dendritic cell costimulator molecule expression is suppressed by T cells in the absence of antigen-specific signalling: role of cluster formation, CD40 and HLA-class II for dendritic cell activation. Immunology 98, 171–180. 10.1046/j.1365-2567.1999.00860.x.

McQuade, A., Coburn, M., Tu, C.H., Hasselmann, J., Davtyan, H., and Blurton-Jones, M. (2018). Development and validation of a simplified method to generate human microglia from pluripotent stem cells. Mol Neurodegener 13, 67. 10.1186/s13024-018-0297-x.

Meleady, L., Towriss, M., Kim, J., Bacarac, V., Dang, V., Rowland, M.E., and Ciernia, A.V. (2023). Histone deacetylase 3 regulates microglial function through histone deacetylation. Epigenetics 18, 2241008. 10.1080/15592294.2023.2241008.

Milacic, M., Beavers, D., Conley, P., Gong, C., Gillespie, M., Griss, J., Haw, R., Jassal, B., Matthews, L., May, B., et al. (2024). The Reactome Pathway Knowledgebase 2024. Nucleic Acids Research 52, D672–D678. 10.1093/nar/gkad1025.

Morabito, S., Miyoshi, E., Michael, N., Shahin, S., Martini, A.C., Head, E., Silva, J., Leavy, K., Perez-Rosendahl, M., and Swarup, V. (2021). Single-nucleus chromatin accessibility and transcriptomic characterization of Alzheimer’s disease. Nat Genet 53, 1143–1155. 10.1038/s41588-021-00894-z.

Muffat, J., Li, Y., Yuan, B., Mitalipova, M., Omer, A., Corcoran, S., Bakiasi, G., Tsai, L.-H., Aubourg, P., Ransohoff, R.M., et al. (2016). Efficient derivation of microglia-like cells from human pluripotent stem cells. Nat Med 22, 1358–1367. 10.1038/nm.4189.

Nam, K.H., and Ordureau, A. (2022). Quantitative proteome remodeling characterization of two human reference pluripotent stem cell lines during neurogenesis and cardiomyogenesis. Proteomics 22, e2100246. 10.1002/pmic.202100246.

Nasser, J., Bergman, D.T., Fulco, C.P., Guckelberger, P., Doughty, B.R., Patwardhan, T.A., Jones, T.R., Nguyen, T.H., Ulirsch, J.C., Lekschas, F., et al. (2021). Genome-wide enhancer maps link risk variants to disease genes. Nature 593, 238–243. 10.1038/s41586-021-03446-x.

Nott, A., Holtman, I.R., Coufal, N.G., Schlachetzki, J.C.M., Yu, M., Hu, R., Han, C.Z., Pena, M., Xiao, J., Wu, Y., et al. (2019). Brain cell type-specific enhancer-promoter interactome maps and disease-risk association. Science 366, 1134–1139. 10.1126/science.aay0793.

Novikova, G., Kapoor, M., Tcw, J., Abud, E.M., Efthymiou, A.G., Chen, S.X., Cheng, H., Fullard, J.F., Bendl, J., Liu, Y., et al. (2021). Integration of Alzheimer’s disease genetics and myeloid genomics identifies disease risk regulatory elements and genes. Nat Commun 12, 1610. 10.1038/s41467-021-21823-y.

Pallotta, M.T., Rossini, S., Suvieri, C., Coletti, A., Orabona, C., Macchiarulo, A., Volpi, C., and Grohmann, U. (2022). Indoleamine 2,3-dioxygenase 1 (IDO1): an up-to-date overview of an eclectic immunoregulatory enzyme. The FEBS Journal 289, 6099–6118. 10.1111/febs.16086.

Panagiotakopoulou, V., Ivanyuk, D., De Cicco, S., Haq, W., Arsić, A., Yu, C., Messelodi, D., Oldrati, M., Schöndorf, D.C., Perez, M.-J., et al. (2020). Interferon-γ signaling synergizes with LRRK2 in neurons and microglia derived from human induced pluripotent stem cells. Nat Commun 11, 5163. 10.1038/s41467-020-18755-4.

Pantazis, C.B., Yang, A., Lara, E., McDonough, J.A., Blauwendraat, C., Peng, L., Oguro, H., Kanaujiya, J., Zou, J., Sebesta, D., et al. (2022). A reference human induced pluripotent stem cell line for large-scale collaborative studies. Cell Stem Cell 29, 1685–1702.e22. 10.1016/j.stem.2022.11.004.

Papageorgiou, I.E., Lewen, A., Galow, L.V., Cesetti, T., Scheffel, J., Regen, T., Hanisch, U.-K., and Kann, O. (2016). TLR4-activated microglia require IFN-γ to induce severe neuronal dysfunction and death in situ. Proc. Natl. Acad. Sci. U.S.A. 113, 212–217. 10.1073/pnas.1513853113.

Patro, R., Duggal, G., Love, M.I., Irizarry, R.A., and Kingsford, C. (2017). Salmon provides fast and bias-aware quantification of transcript expression. Nat Methods 14, 417–419. 10.1038/nmeth.4197.

Prater, K.E., Green, K.J., Mamde, S., Sun, W., Cochoit, A., Smith, C.L., Chiou, K.L., Heath, L., Rose, S.E., Wiley, J., et al. (2023). Human microglia show unique transcriptional changes in Alzheimer’s disease. Nat Aging 3, 894–907. 10.1038/s43587-023-00424-y.

Prinz, M., and Priller, J. (2014). Microglia and brain macrophages in the molecular age: from origin to neuropsychiatric disease. Nat Rev Neurosci 15, 300–312. 10.1038/nrn3722.

Pulido-Salgado, M., Vidal-Taboada, J.M., Barriga, G.G.-D., Solà, C., and Saura, J. (2018). RNA-Seq transcriptomic profiling of primary murine microglia treated with LPS or LPS + IFNγ. Sci Rep 8, 16096. 10.1038/s41598-018-34412-9.

Qiu, S., Sun, M., Xu, Y., and Hu, Y. (2024). Integrating multi-omics data to reveal the effect of genetic variant rs6430538 on Alzheimer’s disease risk. Front Neurosci 18, 1277187. 10.3389/fnins.2024.1277187.

Ramos, D.M., Skarnes, W.C., Singleton, A.B., Cookson, M.R., and Ward, M.E. (2021). Tackling neurodegenerative diseases with genomic engineering: A new stem cell initiative from the NIH. Neuron 109, 1080–1083. 10.1016/j.neuron.2021.03.022.

Rauf, A., Badoni, H., Abu-Izneid, T., Olatunde, A., Rahman, M.M., Painuli, S., Semwal, P., Wilairatana, P., and Mubarak, M.S. (2022). Neuroinflammatory Markers: Key Indicators in the Pathology of Neurodegenerative Diseases. Molecules 27, 3194. 10.3390/molecules27103194.

Roberts, B.S., Anderson, A.G., Partridge, E.C., Cooper, G.M., and Myers, R.M. (2024). Probabilistic association of differentially expressed genes with cis-regulatory elements. Genome Res 34, 620–632. 10.1101/gr.278598.123.

Rogers, B.B., Anderson, A.G., Lauzon, S.N., Davis, M.N., Hauser, R.M., Roberts, S.C., Rodriguez-Nunez, I., Trausch-Lowther, K., Barinaga, E.A., Hall, P.I., et al. (2024). Neuronal MAPT expression is mediated by long-range interactions with cis-regulatory elements. Am J Hum Genet 111, 259–279. 10.1016/j.ajhg.2023.12.015.

Ryan, M., McDonough, J.A., Ward, M.E., Cookson, M.R., Skarnes, W.C., and Merkle, F.T. (2024). Large structural variants in KOLF2.1J are unlikely to compromise neurological disease modeling. Cell Stem Cell 31, 290–291. 10.1016/j.stem.2024.02.006.

Savic, D., Partridge, E.C., Newberry, K.M., Smith, S.B., Meadows, S.K., Roberts, B.S., Mackiewicz, M., Mendenhall, E.M., and Myers, R.M. (2015). CETCh-seq: CRISPR epitope tagging ChIP-seq of DNA-binding proteins. Genome Res 25, 1581–1589. 10.1101/gr.193540.115.

Schoenborn, J.R., and Wilson, C.B. (2007). Regulation of interferon-gamma during innate and adaptive immune responses. Adv Immunol 96, 41–101. 10.1016/S0065-2776(07)96002-2.

Shi, Z., Das, S., Morabito, S., Miyoshi, E., Stocksdale, J., Emerson, N., Srinivasan, S.S., Shahin, A., Rahimzadeh, N., Cao, Z., et al. (2024). Single-nucleus multi-omics identifies shared and distinct pathways in Pick’s and Alzheimer’s disease. 10.1101/2024.09.06.611761.

Siddiqui, S., Liu, F., Kanthasamy, A.G., and McGrail, M. (2024). Stat3 mediates Fyn kinase-driven dopaminergic neurodegeneration and microglia activation. Disease Models & Mechanisms 17, dmm052011. 10.1242/dmm.052011.

Snijders, G.J.L.J., De Paiva Lopes, K., Sneeboer, M.A.M., Muller, B.Z., Gigase, F.A.J., Vialle, R.A., Missall, R., Kubler, R., Raj, T., Humphrey, J., et al. (2023). The human microglia responsome: a resource to better understand microglia states in health and disease. 10.1101/2023.10.12.562067.

Sollis, E., Mosaku, A., Abid, A., Buniello, A., Cerezo, M., Gil, L., Groza, T., Güneş, O., Hall, P., Hayhurst, J., et al. (2023). The NHGRI-EBI GWAS Catalog: knowledgebase and deposition resource. Nucleic Acids Res 51, D977–D985. 10.1093/nar/gkac1010.

Stephens, M. (2016). False discovery rates: a new deal. Biostat kxw041. 10.1093/biostatistics/kxw041.

Stępniak, K., Machnicka, M.A., Mieczkowski, J., Macioszek, A., Wojtaś, B., Gielniewski, B., Poleszak, K., Perycz, M., Król, S.K., Guzik, R., et al. (2021). Mapping chromatin accessibility and active regulatory elements reveals pathological mechanisms in human gliomas. Nat Commun 12, 3621. 10.1038/s41467-021-23922-2.

Sun, E., Motolani, A., Campos, L., and Lu, T. (2022). The Pivotal Role of NF-kB in the Pathogenesis and Therapeutics of Alzheimer’s Disease. Int J Mol Sci 23, 8972. 10.3390/ijms23168972.

Sun, N., Victor, M.B., Park, Y.P., Xiong, X., Scannail, A.N., Leary, N., Prosper, S., Viswanathan, S., Luna, X., Boix, C.A., et al. (2023). Human microglial state dynamics in Alzheimer’s disease progression. Cell 186, 4386–4403.e29. 10.1016/j.cell.2023.08.037.

Taliun, D., Harris, D.N., Kessler, M.D., Carlson, J., Szpiech, Z.A., Torres, R., Taliun, S.A.G., Corvelo, A., Gogarten, S.M., Kang, H.M., et al. (2021). Sequencing of 53,831 diverse genomes from the NHLBI TOPMed Program. Nature 590, 290–299. 10.1038/s41586-021-03205-y.

Tian, M.-L., Ni, X.-N., Li, J.-Q., Tan, C.-C., Cao, X.-P., Tan, L., and Alzheimer’s Disease Neuroimaging Initiative (2019). A Candidate Regulatory Variant at the TREM Gene Cluster Confer Alzheimer’s Disease Risk by Modulating Both Amyloid-β Pathology and Neuronal Degeneration. Front Neurosci 13, 742. 10.3389/fnins.2019.00742.

Trizzino, M., Zucco, A., Deliard, S., Wang, F., Barbieri, E., Veglia, F., Gabrilovich, D., and Gardini, A. (2021). EGR1 is a gatekeeper of inflammatory enhancers in human macrophages. Sci. Adv. 7, eaaz8836. 10.1126/sciadv.aaz8836.

Volpato, V., and Webber, C. (2020). Addressing variability in iPSC-derived models of human disease: guidelines to promote reproducibility. Dis Model Mech 13, dmm042317. 10.1242/dmm.042317.

Wang, L., Zhu, Y., Zhang, N., Xian, Y., Tang, Y., Ye, J., Reza, F., He, G., Wen, X., and Jiang, X. (2024). The multiple roles of interferon regulatory factor family in health and disease. Sig Transduct Target Ther 9, 282. 10.1038/s41392-024-01980-4.

Washer, S.J., Perez-Alcantara, M., Chen, Y., Steer, J., James, W.S., Trynka, G., Bassett, A.R., and Cowley, S.A. (2022). Single-cell transcriptomics defines an improved, validated monoculture protocol for differentiation of human iPSC to microglia. Sci Rep 12, 19454. 10.1038/s41598-022-23477-2.

Xiong, X., James, B.T., Boix, C.A., Park, Y.P., Galani, K., Victor, M.B., Sun, N., Hou, L., Ho, L.-L., Mantero, J., et al. (2023). Epigenomic dissection of Alzheimer’s disease pinpoints causal variants and reveals epigenome erosion. Cell 186, 4422–4437.e21. 10.1016/j.cell.2023.08.040.

Yamanaka, M., Ishikawa, T., Griep, A., Axt, D., Kummer, M.P., and Heneka, M.T. (2012). PPARγ/RXRα-Induced and CD36-Mediated Microglial Amyloid-β Phagocytosis Results in Cognitive Improvement in Amyloid Precursor Protein/Presenilin 1 Mice. J. Neurosci. 32, 17321–17331. 10.1523/JNEUROSCI.1569-12.2012.

Yamasaki, A., Imanishi, I., Tanaka, K., Ohkawa, Y., Tsuda, M., and Masuda, T. (2024). IRF8 and MAFB drive distinct transcriptional machineries in different resident macrophages of the central nervous system. Commun Biol 7, 896. 10.1038/s42003-024-06607-6.

Yan, Z., Gibson, S.A., Buckley, J.A., Qin, H., and Benveniste, E.N. (2018). Role of the JAK/STAT signaling pathway in regulation of innate immunity in neuroinflammatory diseases. Clin Immunol 189, 4–13. 10.1016/j.clim.2016.09.014.

Yang, J., Rajan, S.S., Friedrich, M.J., Lan, G., Zou, X., Ponstingl, H., Garyfallos, D.A., Liu, P., Bradley, A., and Metzakopian, E. (2019). Genome-Scale CRISPRa Screen Identifies Novel Factors for Cellular Reprogramming. Stem Cell Reports 12, 757–771. 10.1016/j.stemcr.2019.02.010.

Yao, L., Berman, B.P., and Farnham, P.J. (2015). Demystifying the secret mission of enhancers: linking distal regulatory elements to target genes. Crit Rev Biochem Mol Biol 50, 550–573. 10.3109/10409238.2015.1087961.

Yaqubi, M., Groh, A.M.R., Dorion, M.-F., Afanasiev, E., Luo, J.X.X., Hashemi, H., Sinha, S., Kieran, N.W., Blain, M., Cui, Q.-L., et al. (2023). Analysis of the microglia transcriptome across the human lifespan using single cell RNA sequencing. J Neuroinflammation 20, 132. 10.1186/s12974-023-02809-7.

Yates, R.L., Pansieri, J., Li, Q., Bell, J.S., Yee, S.A., Palace, J., Esiri, M.M., and DeLuca, G.C. (2022). The influence of HLA-DRB1*15 on the relationship between microglia and neurons in multiple sclerosis normal appearing cortical grey matter. Brain Pathol 32, e13041. 10.1111/bpa.13041.

Yeh, H., and Ikezu, T. (2019). Transcriptional and Epigenetic Regulation of Microglia in Health and Disease. Trends Mol Med 25, 96–111. 10.1016/j.molmed.2018.11.004.

Yeh, F.L., Hansen, D.V., and Sheng, M. (2017). TREM2, Microglia, and Neurodegenerative Diseases. Trends in Molecular Medicine 23, 512–533. 10.1016/j.molmed.2017.03.008.

You, J.-E., Kim, E.-J., Kim, H.W., Kim, J.-S., Kim, K., and Kim, P.-H. (2024). Exploring the Role of Guanylate-Binding Protein-2 in Activated Microglia-Mediated Neuroinflammation and Neuronal Damage. Biomedicines 12, 1130. 10.3390/biomedicines12051130.

Young, A.P., and Denovan-Wright, E.M. (2024). JAK1/2 Regulates Synergy Between Interferon Gamma and Lipopolysaccharides in Microglia. J Neuroimmune Pharmacol 19, 14. 10.1007/s11481-024-10115-z.

Young, A.M.H., Kumasaka, N., Calvert, F., Hammond, T.R., Knights, A., Panousis, N., Park, J.S., Schwartzentruber, J., Liu, J., Kundu, K., et al. (2021). A map of transcriptional heterogeneity and regulatory variation in human microglia. Nat Genet 53, 861–868. 10.1038/s41588-021-00875-2.

Yu, G., and He, Q.-Y. (2016). ReactomePA: an R/Bioconductor package for reactome pathway analysis and visualization. Mol Biosyst 12, 477–479. 10.1039/c5mb00663e.

Yu, G., Wang, L.-G., Han, Y., and He, Q.-Y. (2012). clusterProfiler: an R Package for Comparing Biological Themes Among Gene Clusters. OMICS: A Journal of Integrative Biology 16, 284–287. 10.1089/omi.2011.0118.

Yun, J., Lee, D., Jeong, H., Kim, H.S., Ye, S., and Cho, C. (2021). STAT3 activation in microglia exacerbates hippocampal neuronal apoptosis in diabetic brains. Journal Cellular Physiology 236, 7058–7070. 10.1002/jcp.30373.

Yusa, K., Rad, R., Takeda, J., and Bradley, A. (2009). Generation of transgene-free induced pluripotent mouse stem cells by the piggyBac transposon. Nat Methods 6, 363–369. 10.1038/nmeth.1323.

Zhang, G., Lu, J., Zheng, J., Mei, S., Li, H., Zhang, X., Ping, A., Gao, S., Fang, Y., and Yu, J. (2024). Spi1 regulates the microglial/macrophage inflammatory response via the PI3K/AKT/mTOR signaling pathway after intracerebral hemorrhage. Neural Regen Res 19, 161–170. 10.4103/1673-5374.375343.

Zhao, Z., Liu, A., Citu, C., Enduru, N., Chen, X., Manuel, A., Sinha, T., Gorski, D., Fernandes, B., Yu, M., et al. (2024). Single-nucleus multiomics reveals the disrupted regulatory programs in three brain regions of sporadic early-onset Alzheimer’s disease. Res Sq rs.3.rs-4622123. 10.21203/rs.3.rs-4622123/v1.

