## Supplementary material for "KOLF2.1J iTF-Microglia: A standardized platform to study microglial transcriptional regulatory networks in CNS disease": Document S1

### Supplemental figures

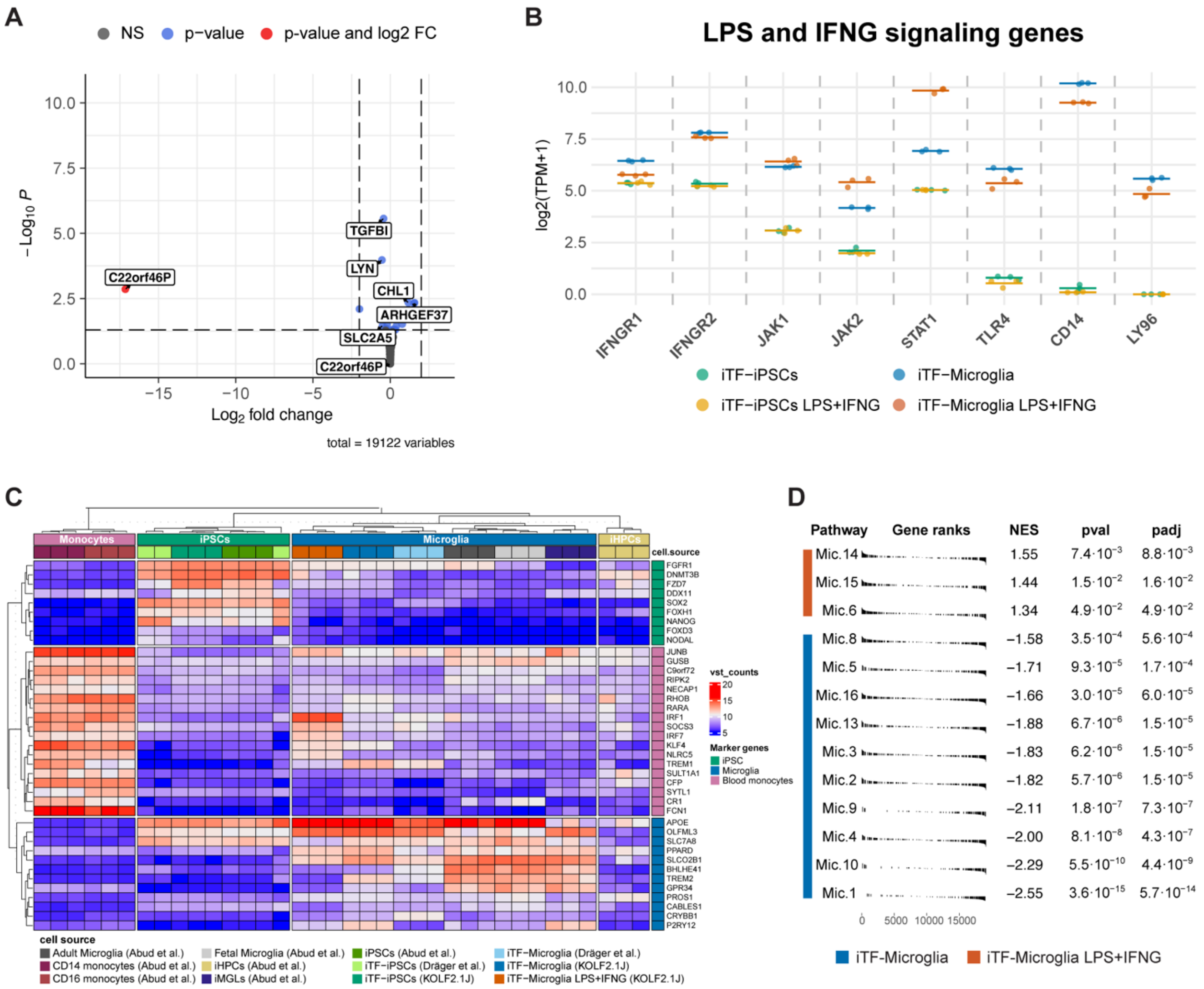


**Figure S1. Transcriptomic analysis of KOLF2.1J iTF-Microglia**

(**A**) Volcano plot of DEGs in LPS+IFNG-treated vs. untreated iTF-iPSCs. Genes with adjusted p-value (padj) < 0.05 and LFC > 2 are considered significant.

(**B**) Expression of IFNG receptors (*IFNGR1*, *IFNGR2*), downstream signaling components (*JAK1*, *JAK2*, *STAT1*), and LPS receptors (*TLR14*, *CD14*, *LY96*). Individual dots represent expression values from each replicate (n = 3). Horizontal bars indicate mean expression across replicates.

(**C**) Hierarchical clustering of KOLF2.1J iTF cells with alongside iPSCs, blood monocytes, and other in vitro-derived myeloid models based on expression of marker genes for pluripotent and myeloid lineages.

(**D**) GSEA of LPS+IFNG-treated vs. untreated iTF-Microglia using gene markers for microglial states. Microglial states include Mic1 (Proliferative), Mic.2-5 (Surveilling), Mic.6-8 (Reacting), Mic.9-10 (Enhanced-redox), Mic.11 (Stress response), Mic.12-13 (Lipid-associated), Mic.14 (Interferon response), Mic15 (Inflammatory) and Mic.16 (SERPINE1 expressing).

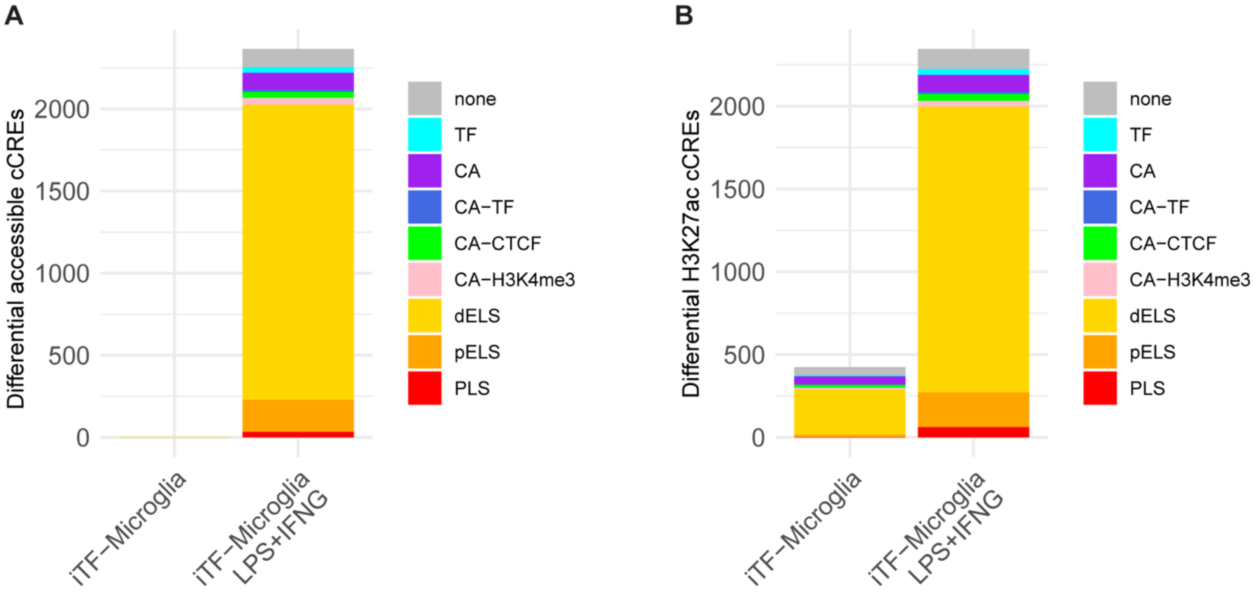


**Figure S2. Chromatin changes in iTF-Microglia following LPS+IFNG stimulation**
(**A**) cCREs with significantly altered chromatin accessibility (FDR < 0.05), annotated using ENCODE4 categories.
(**B**) cCREs with differential H3K27ac enrichment (FDR < 0.05), similarly annotated.
Differential features were identified using csaw analysis.

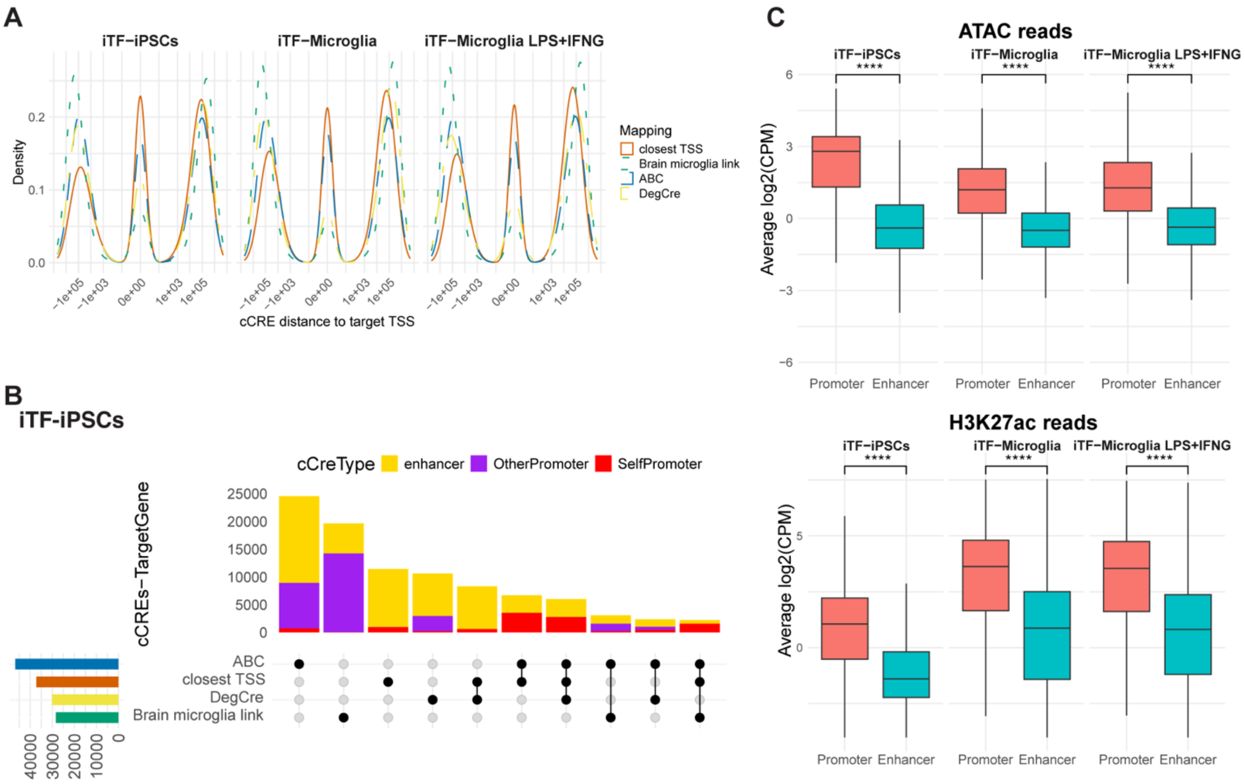


**Figure S3. Properties of cCRE-to-gene associations across mapping models**

(**A**) Distribution of cCRE-to-gene distances for each mapping strategy.

(**B**) Overlap of cCRE-to-gene associations identified by different models in iTF-iPSCs, related to Figure 4A.

(**C**) Boxplots comparing average log2(CPM) reads for ATAC-seq and H3K27ac ChIP-seq signal at promoter- and enhancer-associated cCREs across iTF-iPSCs, iTF-Microglia, and LPS+IFNG-treated iTF-Microglia. Read counts were TMM-normalized, and log2(CPM) values were averaged by group. Statistical comparisons between promoter and enhancer signals were performed using two-sided t-tests (**** $p \leq 0.0001$). Data are represented as boxplots showing the median, interquartile range (IQR), and whiskers extending to 1.5× IQR.

### Supplemental tables

#### Table S8: Description of GWAS datasets, related to Figure 5A and 5B.

| Trait | ID | N | Filename | Reference |
|---|---|---|---|---|
| Alzheimer's disease | (Kunkle) AD | 63926 | Kunkle_etal_Stage1_results.txt | [S1] |
| Alzheimer's disease | (Jansen) AD | 455258 | AD_sumstats_Jansenetal_2019sept.txt.gz | [S2] |
| Alzheimer's disease | (Schwartzentruber) AD | 472868 | GCST90012877_buildGRCh37.tsv.gz | [S3] |
| Alzheimer's disease | (Bellenguez) AD | 788989 | GCST90027158_buildGRCh38.tsv.gz | [S4] |
| Amyotrophic lateral sclerosis | ALS | 138086 | GCST90027164_buildGRCh37.tsv.gz | [S5] |
| Parkinson's disease | PD | 482730 | nallsEtAl2019_excluding23andMe_allVariants.tab.gz | [S6] |
| Multiple sclerosis | MS | 26621 | imsgc_2011_21833088_ms_efo0003885_1_gwas.sumstats.tsv.gz | [S7] |
| Autism spectrum disorder | ASD | 46350 | iPSYCH-PGC_ASD_Nov2017.gz | [S8] |
| Bipolar disorder | BD | 413466 | daner_bip_pgc3_nm_noukbiobank.gz | [S9] |
| Major depressive disorder | MDD | 142646 | daner_pgc_mdd_meta_w2_no23andMe_rmUKBB.gz | [S10] |
| Schizophrenia | SCZ | 130644 | PGC3_SCZ_wave3.european.autosome.public.v3.tsv.gz | [S11] |
| Inflammatory bowel disease | IBD | 34652 | EUR.IBD.gwas_info03_filtered.assoc.gz | [S12] |
| Rheumatoid arthritis | RA | 58284 | RA_GWASmeta_European_v2.txt.gz | [S13] |
| Systemic lupus erythematosus | SLE | 10995 | Meta_Results.zip | [S14] |

**Supplemental reference list**


[S1] Kunkle, B.W., Grenier-Boley, B., Sims, R., Bis, J.C., Damotte, V., Naj, A.C., Boland, A., Vronskaya, M., van der Lee, S.J., Amlie-Wolf, A., et al. (2019). Genetic meta-analysis of diagnosed Alzheimer's disease identifies new risk loci and implicates Aβ, tau, immunity and lipid processing. Nat Genet *51*, 414–430. https://doi.org/10.1038/s41588-019-0358-2.

[S2] Jansen, I.E., Savage, J.E., Watanabe, K., Bryois, J., Williams, D.M., Steinberg, S., Sealock, J., Karlsson, I.K., Hägg, S., Athanasiu, L., et al. (2019). Genome-wide meta-analysis identifies new loci and functional pathways influencing Alzheimer's disease risk. Nat Genet *51*, 404–413. https://doi.org/10.1038/s41588-018-0311-9.

[S3] Schwartzentruber, J., Cooper, S., Liu, J.Z., Barrio-Hernandez, I., Bello, E., Kumasaka, N., Young, A.M.H., Franklin, R.J.M., Johnson, T., Estrada, K., et al. (2021). Genome-wide meta-analysis, fine-mapping and integrative prioritization implicate new Alzheimer's disease risk genes. Nat Genet *53*, 392–402. https://doi.org/10.1038/s41588-020-00776-w.

[S4] Bellenguez, C., Küçükali, F., Jansen, I., Andrade, V., Moreno-Grau, S., Amin, N., Naj, A.C., Grenier-Boley, B., Campos-Martin, R., Holmans, P.A., et al. (2020). New insights on the genetic etiology of Alzheimer's and related dementia (Neurology).

[S5] van Rheenen, W., van der Spek, R.A.A., Bakker, M.K., van Vugt, J.J.F.A., Hop, P.J., Zwamborn, R.A.J., de Klein, N., Westra, H.-J., Bakker, O.B., Deelen, P., et al. (2021). Common and rare variant association analyses in amyotrophic lateral sclerosis identify 15 risk loci with distinct genetic architectures and neuron-specific biology. Nat Genet *53*, 1636–1648. https://doi.org/10.1038/s41588-021-00973-1.

[S6] Nalls, M.A., Blauwendraat, C., Vallerga, C.L., Heilbron, K., Bandres-Ciga, S., Chang, D., Tan, M., Kia, D.A., Noyce, A.J., Xue, A., et al. (2019). Identification of novel risk loci, causal insights, and heritable risk for Parkinson's disease: a meta-analysis of genome-wide association studies. Lancet Neurol *18*, 1091–1102. https://doi.org/10.1016/S1474-4422(19)30320-5.

[S7] International Multiple Sclerosis Genetics Consortium, Wellcome Trust Case Control Consortium 2, Sawcer, S., Hellenthal, G., Pirinen, M., Spencer, C.C.A., Patsopoulos, N.A., Moutsianas, L., Dilthey, A., Su, Z., et al. (2011). Genetic risk and a primary role for cell-mediated immune mechanisms in multiple sclerosis. Nature *476*, 214–219. https://doi.org/10.1038/nature10251.

[S8] Grove, J., Ripke, S., Als, T.D., Mattheisen, M., Walters, R.K., Won, H., Pallesen, J., Agerbo, E., Andreassen, O.A., Anney, R., et al. (2019). Identification of common genetic risk variants for autism spectrum disorder. Nat Genet *51*, 431–444. https://doi.org/10.1038/s41588-019-0344-8.

[S9] Mullins, N., Forstner, A.J., O'Connell, K.S., Coombes, B., Coleman, J.R.I., Qiao, Z., Als, T.D., Bigdeli, T.B., Børte, S., Bryois, J., et al. (2021). Genome-wide association study of more than 40,000 bipolar disorder cases provides new insights into the underlying biology. Nat Genet *53*, 817–829. https://doi.org/10.1038/s41588-021-00857-4.

[S10] Wray, N.R., Ripke, S., Mattheisen, M., Trzaskowski, M., Byrne, E.M., Abdellaoui, A., Adams, M.J., Agerbo, E., Air, T.M., Andlauer, T.M.F., et al. (2018). Genome-wide association analyses identify 44 risk variants and refine the genetic architecture of major depression. Nat Genet *50*, 668–681. https://doi.org/10.1038/s41588-018-0090-3.

[S11] Trubetskoy, V., Pardiñas, A.F., Qi, T., Panagiotaropoulou, G., Awasthi, S., Bigdeli, T.B., Bryois, J., Chen, C.-Y., Dennison, C.A., Hall, L.S., et al. (2022). Mapping genomic loci implicates genes and synaptic biology in schizophrenia. Nature *604*, 502–508. https://doi.org/10.1038/s41586-022-04434-5.

[S12] Liu, J.Z., van Sommeren, S., Huang, H., Ng, S.C., Alberts, R., Takahashi, A., Ripke, S., Lee, J.C., Jostins, L., Shah, T., et al. (2015). Association analyses identify 38 susceptibility loci for inflammatory bowel disease and highlight shared genetic risk across populations. Nat Genet *47*, 979–986. https://doi.org/10.1038/ng.3359.

[S13] Okada, Y., Wu, D., Trynka, G., Raj, T., Terao, C., Ikari, K., Kochi, Y., Ohmura, K., Suzuki, A., Yoshida, S., et al. (2014). Genetics of rheumatoid arthritis contributes to biology and drug discovery. Nature *506*, 376–381. https://doi.org/10.1038/nature12873.

[S14] Julià, A., López-Longo, F.J., Pérez Venegas, J.J., Bonàs-Guarch, S., Olivé, À., Andreu, J.L., Aguirre-Zamorano, M.Á., Vela, P., Nolla, J.M., de la Fuente, J.L.M., et al. (2018). Genome-wide association study meta-analysis identifies five new loci for systemic lupus erythematosus. Arthritis Res Ther *20*, 100. https://doi.org/10.1186/s13075-018-1604-1.
